# Microbial community dynamics and coexistence in a sulfide-driven phototrophic bloom

**DOI:** 10.1101/604504

**Authors:** Srijak Bhatnagar, Elise S. Cowley, Sebastian H. Kopf, Sherlynette Pérez Castro, Sean Kearney, Scott C. Dawson, Kurt Hanselmann, S. Emil Ruff

**Author notes:** Denotes equal contribution (Srijak Bhatnagar, Elise S. Cowley).

## Abstract

Phototrophic microbial mats commonly contain multiple phototrophic lineages that coexist based on their light, oxygen and nutrient preferences. Here we show that similar coexistence patterns and ecological niches can occur in suspended phototrophic blooms of an organic-rich estuary. The water column showed steep gradients of oxygen, pH, sulfate, sulfide, and salinity. The upper part of the bloom was dominated by aerobic phototrophic *Cyanobacteria*, the middle and lower parts were dominated by anoxygenic purple sulfur bacteria (*Chromatiales*) and green sulfur bacteria (*Chlorobiales*), respectively. We found multiple uncultured phototrophic lineages and present metagenome-assembled genomes of two uncultured organisms within the *Chlorobiales*. Apparently, those *Chlorobiales* populations were affected by *Microviridae* viruses. We suggest a sulfur cycle within the bloom in which elemental sulfur produced by phototrophs is reduced to sulfide by *Desulfuromonas sp*. These findings improve our understanding of the ecology and ecophysiology of phototrophic blooms and their impact on biogeochemical cycles.

## Background

Estuarine and coastal water bodies are dynamic and ubiquitous ecosystems that are often characterized by the mixing of terrestrial freshwater and ocean saltwater. Brackish habitats can have striking physical and chemical characteristics that differ from both fresh and saltwater ecosystems [1, 2]. Brackish ecosystems are very diverse and support large microbial and macrobial communities [1]. Estuaries also provide crucial ecosystem services, the most salient of which are trapping and filtering terrestrial runoffs and pollutants before they enter the oceans, coastal protection, erosion control and habitat-fishery linkages [3–6].

Estuaries harbor abundant and diverse microbial communities that form the basis of a complex food chain by fixing carbon dioxide through photosynthesis or chemosynthesis [7–9]. Additionally, carbon introduced to estuaries as organic matter from the oceans or land can be remineralized by heterotrophic microbial communities [10–12]. The decomposition of sulfur containing organic compounds through fermentation can lead to the production of sulfide in estuarine sediments [13]. Furthermore, sulfate brought into estuaries by ocean waters can be used by sulfate reducers, which reduce sulfate into elemental sulfur or sulfide [13, 14]. Sulfate introduced by ocean water and sulfide released from the sediments form gradients in the water column that cause a chemocline in the brackish water column [15]. Additionally, estuaries and coastal marshes often exhibit a halocline, i.e. a change in salinity, and the depletion of oxygen in the water column can create an oxycline [16, 17]. Overlapping gradients, e.g. in salinity, light availability, as well as oxygen and sulfide concentration can create habitats and niches that favor certain microbial communities and conversely microbial communities can affect and respond to such gradients [18–20].

Gradients of light availability, oxygen and sulfur concentrations in stratified aquatic environments offer the development for complex and stable microbial assemblages [21]. These gradients are usually divided into a surface layer rich in oxygen, an intermediate layer with decreasing oxygen and a bottom anoxic layer. The surface layer is often dominated by oxygenic phototrophic microorganism such as *Cyanobacteria*. The anoxic layer, particularly in systems with high organic loads or seagrass beds, provides niches for anaerobes such as sulfate-reducing bacteria [22]. In the intermediate layer, anoxygenic phototrophs use the light from the surface and the sulfide from the bottom layers [23]. The biogeochemical processes leading to stratification in phototrophic blooms are relatively well understood [24], yet ecological niches, microbial interactions and community dynamics are less well constrained.

The abiotic and biotic drivers of stratified aquatic environments such as estuaries can fluctuate frequently and rapidly as a result of many factors, for example, tidal cycles, weather patterns, and seasonal cycles [25–30]. Such disturbances can bring about noticeable changes in the microbial community structure of the ecosystem. It was shown that estuarine communities were structured by salinity [31–34], precipitation [32, 35], temperature [33, 34], oxygen [35, 36] and also seasonal changes [34]. These community shifts included changes in phytoplankton populations with salinity [31], declining populations of *Rhodobacterales* with decreasing salinity [35], declining populations of phototrophic *Ca.* Aquiluna with decreasing oxygen concentration, as well as general changes in the richness and evenness of the community [31–36].

Trunk River in Woods Hole, MA is a brackish ecosystem, on the coast of Vineyard Sound (N 41.535236, W −70.641298). Near the mouth, Trunk River forms a shallow lagoon where freshwater mixes with seawater. Storms, tides, and run-off introduce large amounts of biomass to the pond forming thick layers of decaying seagrass and other organic matter. The pond has a distinct sulfidic odor that emanates from the water and gases bubble up from the sediment. Bright yellow microbial blooms can be observed occasionally just below the water surface (see Figure 1, S1), forming and disappearing within days to weeks. Transient blooms were observed to occur in natural depressions in the decaying organic matter and anecdotally seemed to be initiated by physical disturbance events, potentially from storms, tidal extremes, human activity, or animals. Given this visually striking natural ecological progression, we set out to test whether physical disturbance could trigger the blooms, and whether the established blooms could be used as a model system to investigate the microbial ecology and ecophysiology of sulfur-oxidizing phototrophs.

**Figure 1:**
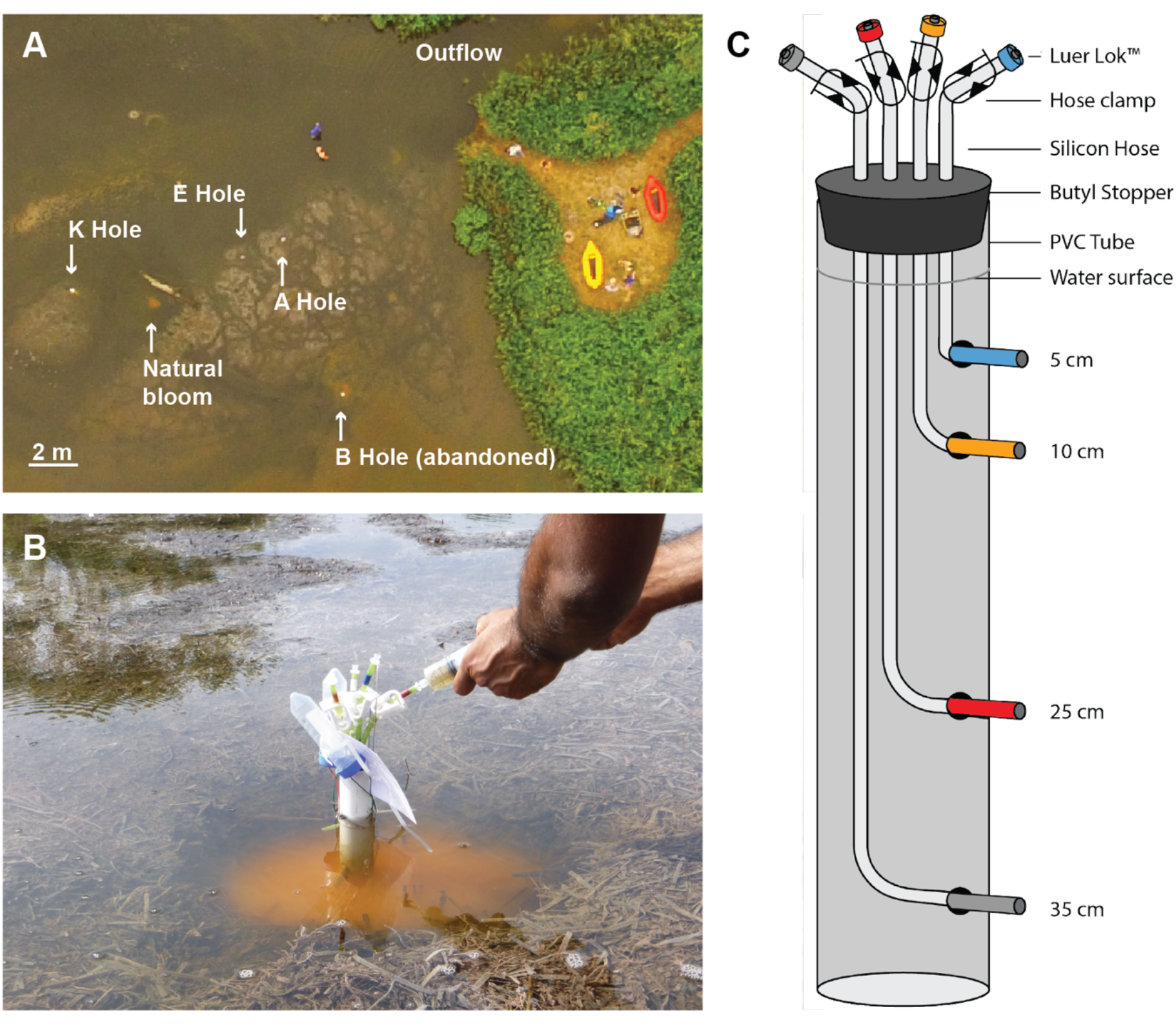
Sampling sites. **A.** Aerial view of experimental sites (A, E, and K) in the Trunk river lagoon. The water enters the lagoon from the right and exits to the sea through a channel marked outlet. **B.** View of a **LEMON** (**L**ong-term **E**nvironmental **MON**itoring) sampling device during sample collection on time point 3; 5 days post-disturbance. **C.** Schematic of the LEMON.

We mimicked physical disturbances of the brackish ecosystem by creating artificial depressions in the decaying organic matter, and monitored the microbial community response and population dynamics, as well as investigated ecological niches of the key populations. Based on the above described observations of thick layers of decaying organic matter and naturally occurring, rapid blooms of phototrophs, we hypothesized that i) the disturbance would release sulfide from the sediment and cause a sulfide-driven phototrophic bloom ii) due to its rapid development the bloom would likely be dominated by very few populations and iii) steep physicochemical gradients would establish that create (transient) anoxic habitats in the water column analogous to blooms in stratified lakes. We discuss the resulting reproducible ecological succession and provide insights into the habitats, niches and coexistence of phototrophic microorganisms. Our findings contribute to the understanding of the ecological processes and dynamics that shape phototrophic blooms, which are a naturally occurring phenomenon in many ecosystems.

## Results

This study was designed to investigate microbial community assembly, community turnover and syntrophic interactions in a sulfide-driven phototrophic bloom. To gain insights into the microorganisms niches and potential key metabolisms we studied the physicochemistry of the water column and the diversity of photopigments, and performed amplicon and metagenomic sequencing.

### Physicochemistry of the water column

At the first sampling time point (2 days post-disturbance), no difference in color was observed in the water column. Two days later, a faint pink layer was observed in the water column, and faint shades of yellow appeared in samples from the 25 cm depth (Figure S2, Supplementing Results). The yellow color of the suspension was most intense from timepoint 4 to 7 and had almost disappeared by timepoint 8. Within the first three days of the experiment the pH decreased between one and two units in all layers, with lowest values present in the deepest layer (Figure 2). Over the 15-day sampling period, pH showed more variation in the two upper layers than in the two deeper layers where it was very constant at values between 6 and 6.3. Throughout the experiment the water column in all three experiments showed a stable halocline with brackish water (5 ‰ salinity) at the water surface and saltwater (30 ‰) at 35 cm depth (Figure 2). Salinity increased with depth and was 12 ‰ and 23 ‰ at 10 cm and 25 cm, respectively. Major ions also reflect this trend (e.g. calcium, potassium in Figure S6). The dissolved oxygen (DO) concentrations showed a relatively stable oxycline between 10 and 25 cm. At 10 cm and above, DO was mostly higher than 50 µM (91±45 µM) corresponding to ∼20 % oxygen saturation (36±17 %). At 25 and 35 cm DO was mostly below 50 µM (23±18 µM), hence below ∼20 % (9±9 %) saturation. The oxygen concentration slowly decreased in the upper two layers during the first half of the experiment but recovered again to the original values towards the end of the experiment. At 5 and 10 cm, DO averaged over the experiment was 101±47 µM and 81±41 µM, respectively (Figure 2). At 25 and 35 cm, the average DO was 28±22 µM and 17±11 µM, respectively. The sulfate concentrations in the water column decreased along the depth gradient, with the highest sulfate concentration at 5 cm (≈ 2 mM) and the lowest at 25 cm (≈ 0.2 mM) (Figure 2). In contrast, the sulfide concentrations were lowest at 5 cm (Figure 2F). Interestingly, the greatest sulfide concentration was measured at 10 cm depth peaking at over 1 mM towards the end of the experiment. Below 10 cm, sulfide concentration was still high, but declined to 0.75 mM ± 0.22 at 25 cm and 0.5 mM ± 0.17 at 35 cm. The normalized biomass measured for the 5 cm samples throughout the sampling period was nearly zero (Figure 2). At 10 cm, 25 cm, and 35 cm, the normalized biomass measured was approximately, 0.2 mg/mL, 0.3 mg/mL, and 0.2 mg/mL, respectively. For details concerning iron (Fe(II), Fe(III), total Fe), nitrate, calcium, potassium, ammonium and acetate refer to Supplementary Results and Figure S6. Overall, the measurements revealed stable and reproducible physicochemical gradients that divided the previously homogenous water column into layers with different redox conditions and energy availability.

**Figure 2:**
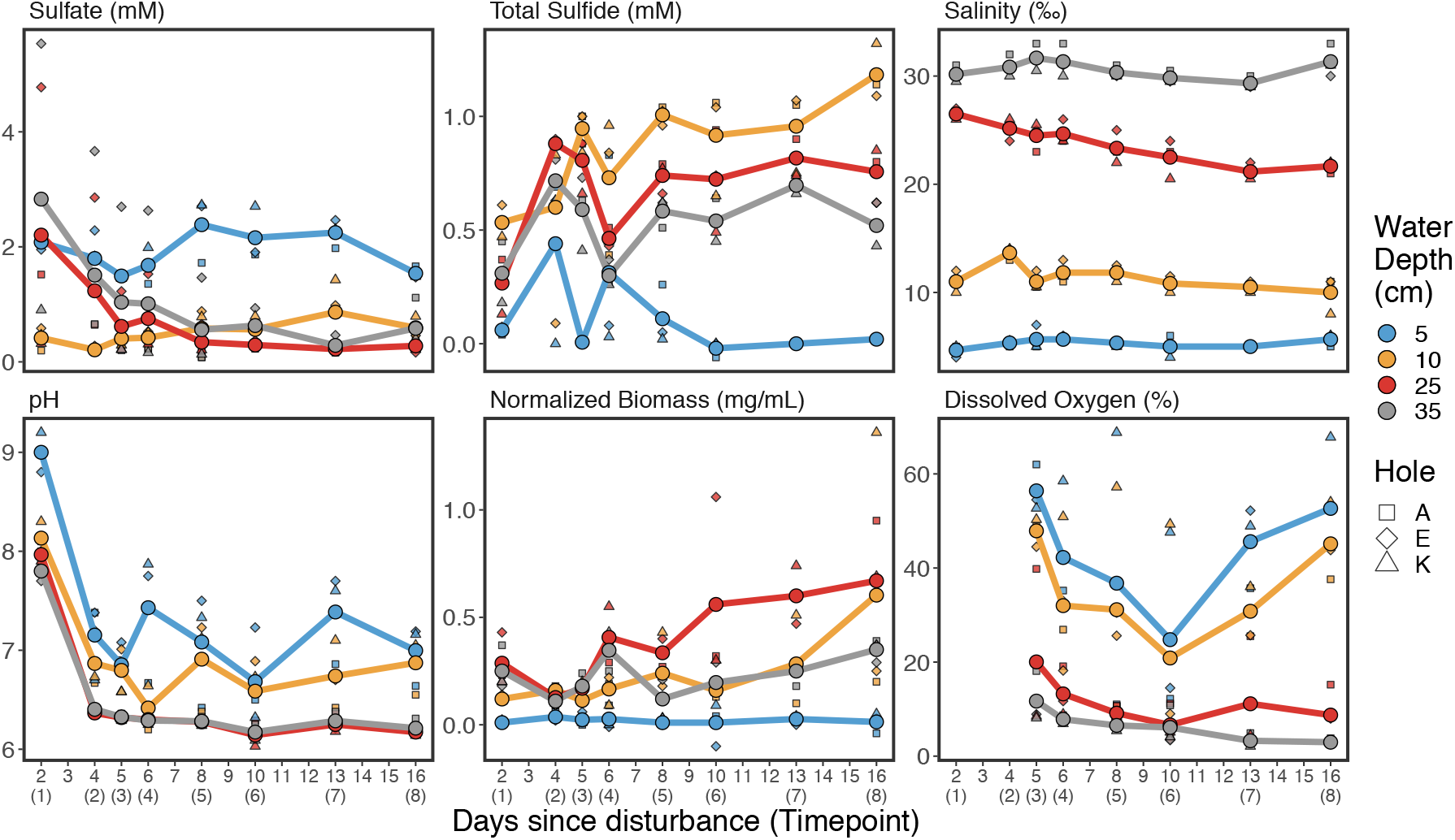
Physicochemical measurements at the sampling sites. Measurements are shown as averages (circles) across the three replicate holes. Measurements at individual holes are shown as squares, diamonds and triangles, the trend is shown as lines connecting average values. The x-axis shows days since disturbance and sampling timepoints in brackets. The y-axis shows the respective units. For an alternative representation of the physicochemical parameters as depth profiles instead of temporal profiles, see Figure S5. For further parameters (Fe (II); Fe (III); Total Fe, nitrate) refer to Figure S6.

### Spectral Absorbance of Phototrophic Community

We measured absorbance spectra from filters of samples from A, E and K experiments (Figure 5A) and compared the spectra to those of representative cultured species of the most abundant phototrophic genera from the literature [37–41] (Figure 5B). Our results show that pigments belonging to PSB (purple vertical bands) were prominent in the upper layer of the bloom (orange spectra in 5A) especially between day 10 and 13, while GSB pigments (green vertical band in 5A) dominated the lower layers of the bloom (red and gray spectra) starting at day 10. Pigments characteristic for *Cyanobacteria* (brown vertical band) were less abundant in the bloom but increased at the end of the experiment relative to the PSB and GSB peaks. This suggests a minor role of *Cyanobacteria* initially and during the bloom but a more important role upon return to equilibrium. Pigments of all major phototrophs were detected throughout the experiment (gray vertical band). The spectral results suggest the coexistence of multiple phototrophic lineages over the entire duration of the experiment.

### Microbial Community Structure and Taxonomic Composition

At the beginning of the experiment, the microbial diversity was high in all four water depths and very similar across replicate ecosystems. Alpha diversity rapidly decreased with the onset of the bloom, and within two days the communities in the four depth layers substantially changed (Figure 4, 5, S7, S8). The bloom occurred between 10 – 30 cm water depth (Figure S2) with highest cell numbers (peaking at >10^8^ cells ml^-1^) and biomass at around 25 cm water depth (Figure 2, S4) in brackish, mildly acidic, and hypoxic waters (Figure 2). The number of observed amplicon sequence variants (ASV), as well as estimated richness, Shannon entropy, and Inverse Simpson diversity significantly decreased between the surface water and the water at a depth of 10 cm and 25 cm (Figure 4; p=0.001). This change is most striking in the case of Inverse Simpson diversity, a measure for evenness. In just one day, evenness dropped in both 10 cm and 25 cm water depth by over one order of magnitude to low single digit values (Figure 4; Table S1). This means the community was dominated by one ASV (a pure culture has an Inverse Simpson diversity index of 1). This decrease in diversity was accompanied by a substantial decrease in pH, as well as an increase in sulfide concentration.

The rapid and profound change of community structure is corroborated by a high turnover of ASV between the layers and timepoints (Figure 5, S8). The top layer is well separated from the deeper layers. The communities at 25 cm water depth experienced the largest turnover, i.e. change in community structure, and showed a loss in diversity during the experiment that seemed to have recovered at the last time point (Figure 5). The communities of all three deep layers (15 – 35 cm) had a similar community structure at the beginning of the experiment. Interestingly, during the course of the experiment the community structure of each layer followed a different trajectory, yet at the end converged again. The trajectories of layer 2-4 indicate that the bloom shifted the microbial communities in these layers to an alternative stable state.

The taxonomic composition was assessed at all phylogenetic levels (Figure 6, S9). We observed a total of 73 bacterial phyla. The surface community (5 cm) remained relatively unchanged throughout the experiment and was dominated by *Proteobacteria*, *Chlorobi*, *Cyanobacteria* and *Actinobacteria*. The communities in the deeper oxygen poor and sulfide rich zones (10 – 35 cm) were more dynamic, being dominated by *Bacteroidetes*, *Proteobacteria*, *Firmicutes*, and *Chloroflexi*. In general, taxonomic diversity was highest in the deepest layer (35 cm). The profound change in diversity was accompanied by a change in composition. Within a few days, there was a substantial increase in the abundance of *Chlorobi*, which comprised more than 75% of the community at that time. This increase persisted for nine days, but levelled off at the end of the experiment. The datasets of all layers and timepoints were dominated by ASVs affiliating with phototrophic organisms, as shown by relative sequence abundances on genus level (Figure 6A). Some phototrophs occurred in all layers at similar relative sequence abundances, such as *Halochromatium* and “*Candidatus* Chloroploca”. The stable surface layer harbored *Cyanobium* and “*Candidatus* Aquiluna”, which decreased in the deeper layers. The upper layer of the bloom showed an increased relative sequence abundance of *Allochromatium,* the lower bloom layer was dominated by *Prosthecochloris* and *Chlorobaculum* (Figure S11). In addition to phototrophs the bloom layers were enriched with sulfur-reducing *Desulfuromonas sp.* as well as *Exiguobacterium sp.* (Figure 6, Figure S9, S11). The layer above the bloom was slightly enriched with sulfur-oxidizing *Thiovirga sp.* and the layer below the bloom with *Erypsipelothrix sp.* Sulfate-reducing *Desulfobacteraceae* and *Desulfobulbaceae* were observed at low relative abundances in all layers (Figure S9B).

Interestingly, almost all *Prosthecochloris* affiliated reads belonged to a single sequence variant, while ASV diversity affiliated with the closely related *Chlorobaculum* increased over time (Figure 6B, S10). The relative sequence abundance of *Chlorobiales* noticeably changed at 25 cm depth, where the yellow microbial bloom was visually observed. *Chlorobiales* ASVs accounted for >25 % of reads in our dataset. To identify the phylogeny of ASV affiliating with *Chlorobiales*, we placed the representative sequence of each ASV on a reference tree of known *Chlorobiales*. The most abundant *Chlorobiales* ASV (ASV_1) affiliated with the genus *Prosthecochloris*, and specifically in the monophyletic clade of *Prosthecochloris vibrioformis* (Figure S12), followed by an ASV (ASV_2) affiliating with *Chlorobaculum*. Together, these two ASVs account for >97 % of the *Chlorobiales* reads. In general, we found a high number of unclassified lineages. The 20 most abundant ASV accounted for about 50 % of all sequences, twelve of those belonged to unclassified genera or families (Figure S9B). The novelty was especially high within the *Chromatiaceae* five of the eight ASV that ranked among the “top 20” belonged to an unclassified genus.

### Metagenomics-derived insights into *Chlorobiales* populations

We calculated the index of replication (iRep) [42] of the *Prosthecochloris* and *Chlorobaculum* populations based on the metagenome-assembled genomes (MAGs) that were recovered from the community metagenomes of two replicate experiments (Replicates A, E) and the enrichment culture (SK) at timepoint 7. Both populations were replicating rapidly. *Prosthecochloris* (bin10) had an iRep value of 3.7 (r^2^=0.90, sample 7A3), which indicates that on average every cell had 2.5 replication events at the time of sampling. *Chlorobaculum* (bin 6) had iRep values of 2.5 (r^2^=0.95, sample 7E3) and 2.8 (r^2^=0.95, sample 7K3), indicating that on average every cell had ∼1.5 replication events. Both MAGs contained genes of the thiosulfate-oxidizing multienzyme system (Sox). Bin 6 contained SoxXYZAB while Bin 10 contained only SoxYZ (Figure S17). Both MAGs also contained the sulfate reductase subunit DsrAB involved in sulfate reduction, and components of assimilatory sulfate reduction (CysND and Cys). Genes for bacteriochlorophhyll (BChl) biosynthesis (bchEMU) were found in both MAGs. Bd-type oxidases (CydAB) were present in both MAGS, while heme-copper oxygen reductases were only found in Bin 6 including several cytochrome c oxidases (COX10, CyoABCDE and III) (Table S4).

Bin 6 (*Chlorobaculum sp.*) and bin 10 (*Prosthecochloris sp.*) contain CRISPR arrays denoted as either type I (cas3) or III (cas10) CRISPR systems [43] (Figure S18, S19). CRISPR predictions revealed 3 direct repeat sequences in both MAGs of 30 and 35 (2) bp in length for Bin 6 and 37, 32, and 33 for the Bin 10 (Table S5). None of the spacers were shared by the closest reference and representative genomes or matched sequences in the CRISPR database [44]. However, a highly similar CRISPR array and direct repeat sequence were found between our Bin 6 and *Chlorobaculum parvum* NCBI8327 with 60% *cas* genes similarity (Figure S18). The metagenomes of all experiments, as well as of the GSB enrichment culture contained high relative sequence abundances of viruses affiliating with *Microviridae* (Figure S20).

## Discussion

In this study, we created artificial depressions in the decaying seagrass layer of Trunk River to mimick disturbances that naturally occur at this site. We performed triplicate experiments that showed very similar physicochemical gradients and patterns of community structure enabling us to study microbial community succession in a natural setting. The observed slight variations between replicate sites were likely due to differences in the organic matter composition and distance to the lagoon access, or due to ripples and disturbances caused by weather, animals, and sampling scientists. The replicated disturbances of the organic matter layers (A-hole, E-hole, and K-hole) released trapped sulfide and caused the rapid establishment of steep physicochemical gradients as well as the development of a bloom of sulfide-oxidizing phototrophs. We monitored the microbial community assembly and succession, highlight the ecological niches of key populations and indicate syntrophic interactions between phototrophs and sulfur reducers.

### Sulfur cycling in the phototrophic bloom

Sulfate concentrations in the bottom layers decreased substantially within the first days and were lowest in the bloom layer at 25 cm depth, where sulfate was almost entirely depleted. We found sulfate-reducers affiliating with *Desulfobacteraceae* and *Desulfobulbaceae* in the hypoxic layers of the bloom (Figure S9B) likely producing sulfide using either hydrogen or organic acids, e.g. acetate (Figure S6) released from fermented organic matter. The sulfide concentrations were highest at the upper boundary of the bloom at 10 cm water depth after the system stabilized around day 6 (Figure 2). This is unexpected since reduced sulfur species, especially hydrogen sulfide, are the electron donor for the green and purple phototrophs and thus should have been depleted in these layers. At the same time, we found an increased relative abundance of sulfur-reducing *Desulfuromonas sp.* in the bloom layers, peaking at around 15 % relative sequence abundance. *Desulfuromonas sp.* are known to live in freshwater ecosystems and reduce elemental sulfur to sulfide [45–47], which in turn can be reused by the sulfide-oxidizing phototrophs. Our findings suggest that the initially present sulfide was released from the sediment but was likely replenished by sulfate reducers from sulfate, as well as by sulfur reducers from sulfur. Sulfide (and thiosulfate) are oxidized to elemental sulfur by the anoxygenic phototrophs and hence the potential sulfur reduction by *Desulfuromonas sp.* indicates sulfur cycling carried out by multiple species (Figure 7). A similar synergistic interaction was suggested to occur in Lake Cadagno between sulfur disproportionating *Desulfocapsa thiozymogenes* and purple sulfur bacteria affiliating with *Lamprocystis* [48]. At early timepoints the microbial suspension was beige and milky, indicating the presence of large amounts of elemental sulfur in the sample (Figure S2). Later the samples turned more yellow, due to an increase in phototrophic organisms and their photopigments (Figure 2, 3), but also cleared up and became translucent (Figure S2). Taken together these findings could mean that *Desulfuromonas sp.* reduced the elemental sulfur (possibly present as polysulfides) that was produced by the anoxygenic phototrophs before it accumulated in the suspension. An observation that merits future research. Such a sulfur loop provides a positive feedback that could explain the enrichment of sulfide in the bloom as well as the very rapid development of the sulfur-oxidizing phototrophs. The involved *Chlorobi* and *Deltaproteobacteria* could even form tight aggregates similar to *Chlorochromatium aggregatum* [49], to efficiently use the sulfur intermediate.

**Figure 3:**
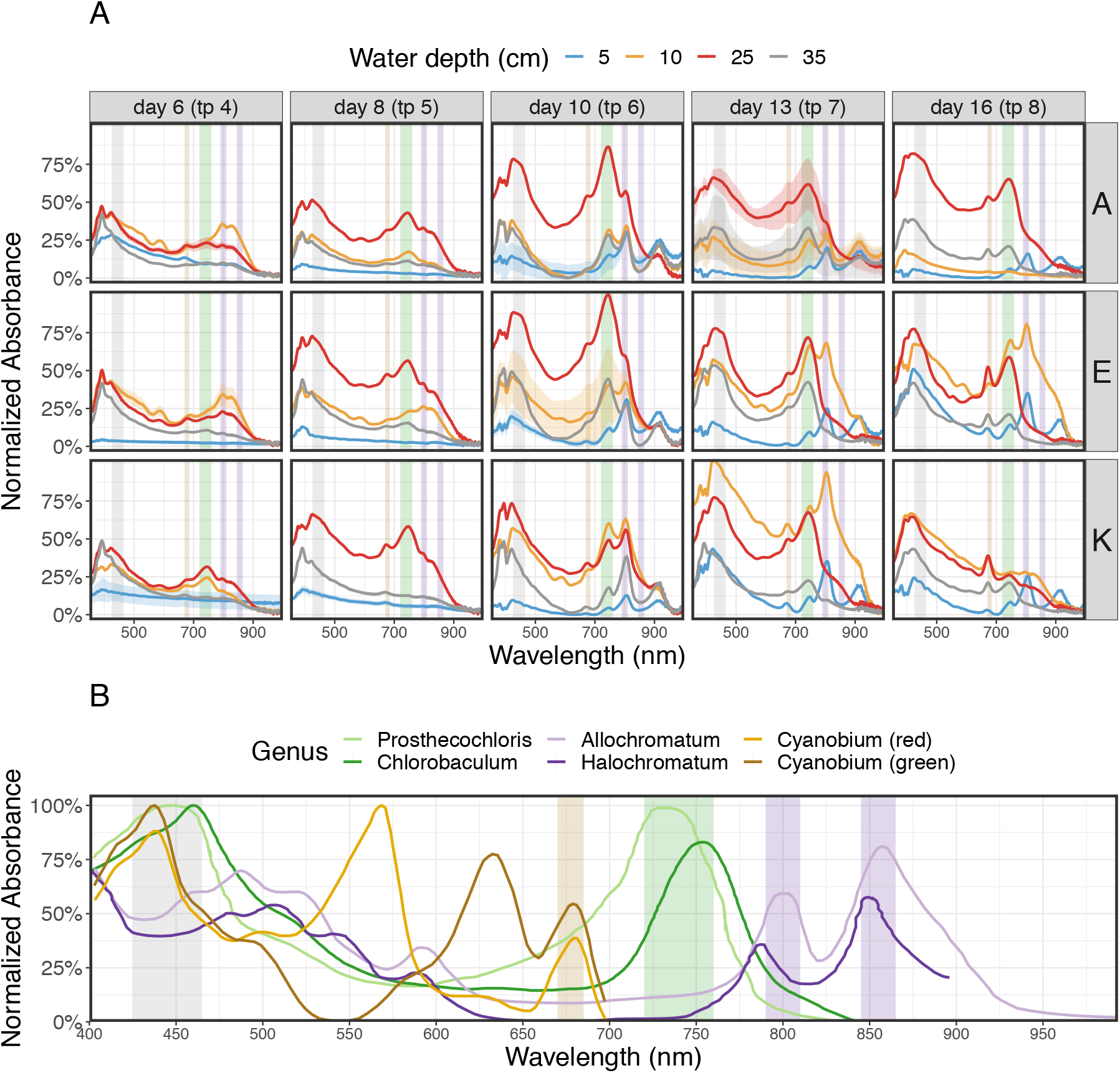
Spectral Absorbance. A) Sample spectra for the 3 sites at 5 different time points each showing averages of at least three replicate spectral analyses per sample. Significance bands along the spectra indicate one standard deviation (bands are mostly smaller than center line and thus not visible). Vertical bands indicate major absorbance peaks of the GSB group in green (*Prosthecochloris* and *Chlorobaculum*, 720-760 nm) and the PSB group purple (*Allochromatium* and *Halochromatium*, 790-810 nm and 845-865 nm) highlighting the transient appearance and likely dominance of these phototrophs over the course of the experiments. Also indicated is the general phototroph absorbance band in the 425-465 nm window in light gray. Cyanobacterial groups have distinct absorbance peaks in the 500-700 nm range that are not prominent in the sample spectra except for the characteristic 670-685 nm peak (light brown vertical band) reflecting the presence but likely smaller role of these taxa during the experiment. B) Absorbance spectra of pure culture representatives of major phototrophic groups with the same vertical bands as in panel A highlighting diagnostic absorbance peaks of GSB (in green), PSB (in purple), cyanos (in orange/brown) and phototrophs in general (in gray). All absorbance spectra were normalized to the respective highest peak.

### Assembly and coexistence of phototrophic microorganisms

The multispecies phototrophic bloom (fondly termed “microbial lemonade”, Figure 1C) formed around two to four days post-disturbance and was fully established by day six. The bloom contained lineages from multiple phyla but was dominated by green and purple sulfur bacteria. The color of the bloom slightly shifted from yellow-white at early timepoints to yellow-orange at mid timepoints to yellow-green at late timepoints (Figure S2), likely due to the relative influence of the photopigments of green and purple sulfur bacteria. The change in bacteriochlorophylls is reflected by the pigment spectra collected at the different timepoints (Figure 3). The opacity and color of the suspension, especially at the beginning of the experiment, is likely influenced by the presence of polysulfides that are produced abiotically [50], as well as biotically by purple and green sulfur bacteria due to their lack of the soxCD genes [51].

Interestingly, the sequencing data suggested that especially the lower layer of the bloom was dominated by an apparently clonal population of green sulfur bacteria affiliated with *Prosthecochloris vibrioformis*. The green sulfur bacteria are sulfur-oxidizing, strictly anaerobic, obligate photoautotrophs [52]. Yet, based on our oxygen measurements, the Trunk River GSB populations apparently tolerated relatively high oxygen concentrations of around 30 µM, but up to 80 µM (Figure 2). The low concentration of dissolved oxygen at 25 cm depth combined with sulfide, salinity, and low light created an optimal habitat for *Prosthecochloris sp*. The observed community turnover (Figure 4) indicates that communities in the layers 2-4 shifted from one stable state at the beginning of the experiment (timepoint 1) to an alternative stable state at the end of the experiment (timepoint 8). It appears that PSB (*Allochromatium sp.*) played a key role in stable state one, while the community of stable state two was equally dominated by both GSB populations (*Prosthecochloris sp.* and *Chlorobium sp.*). The change of relative abundances of phototrophs over the course of the experiment seems to be responsible for the pronounced community turnover, because together these few clades made up the majority of sequence reads. The high affinity for sulfide and the efficient growth of GSB at low light conditions [53] may have enabled GSB to outcompete PSBs at the end of the experiment leading to a community adapted to the changed conditions.

**Figure 4:**
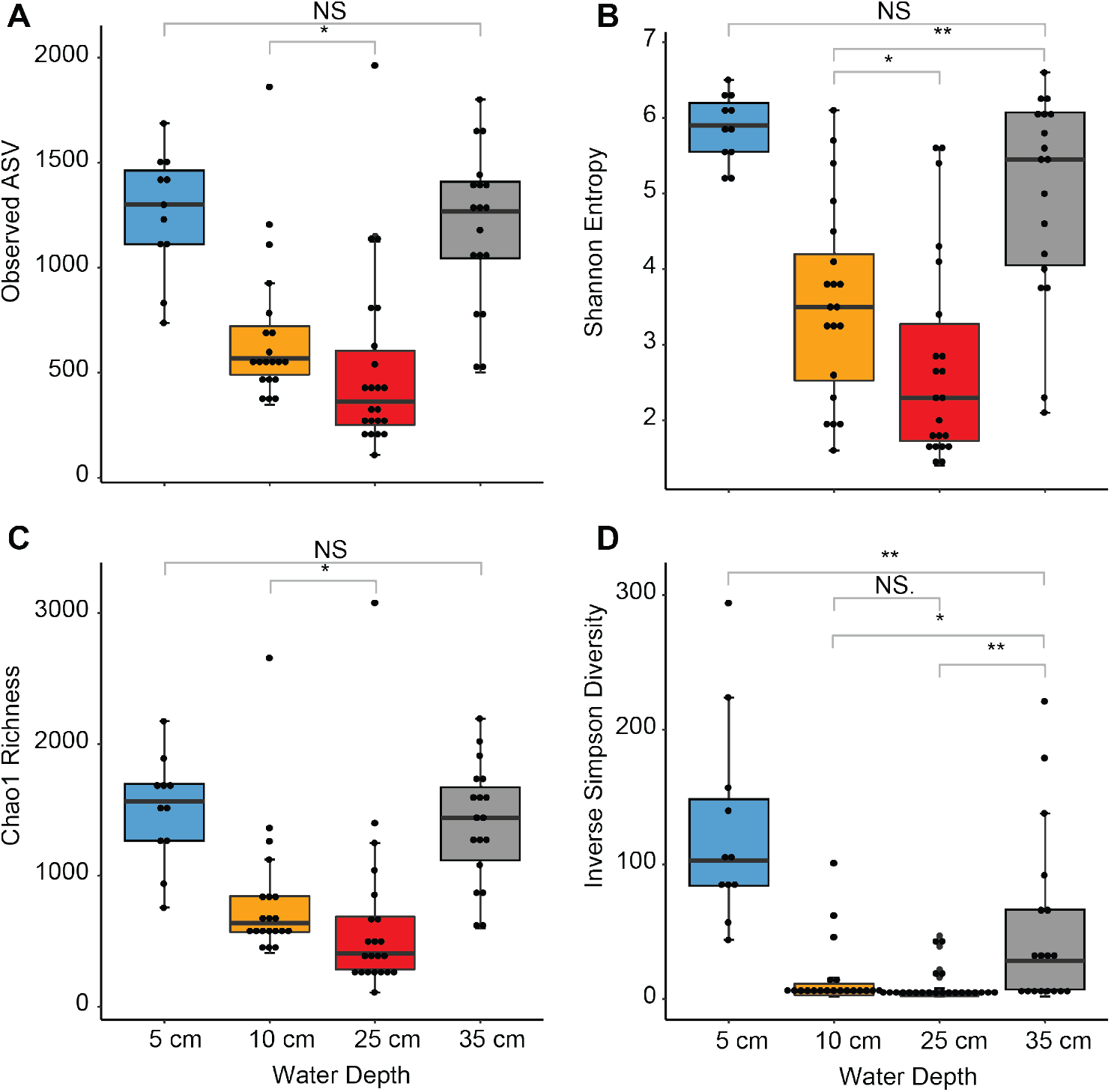
Diversity Indices of all samples shown as boxplots grouped by depth. Pairwise comparisons with low significance levels are shown (NS, *: p<0.1, **: p<0.01). All pairwise comparisons that are not shown were highly significant (***: p<0.001), e.g. panel A 5 cm vs 10 cm.

Despite the dominance of few populations the disturbance created a habitat with gradients of pH, salinity, light, oxygen, and sulfide that enabled the coexistence of multiple phototrophic clades from at least five different phyla (*Actinobacteria, Chlorobi, Chloroflexi, Cyanobacteria and Gammaproteobacteria*). Coexistence of multiple phototrophic lineages was observed before, especially in lakes [21,54,55]. The coexistence of organisms competing for the same energy source is due to the different absorption maxima of each clades’ photopigments (Figure 3), as well as their need for different electron donors, and the varying salinity and oxygen tolerances of each clade. At Trunk River *P. vibrioformis* is absent at 5 cm and present only in low abundance at 10 cm. The surface layer (5 cm depth) is inhabited by oxygenic phototrophic *Cyanobacteria* affiliating with *Cyanobium*, while the upper layer of the bloom (10 cm depth) is dominated by purple sulfur bacteria of the order *Chromatiales* (Figure 6). Because *Prosthecochloris* are adapted to low light conditions [56] and respond to different wavelengths of light than *Cyanobacteria* and photosynthetic *Proteobacteria* [57, 58], it is reasonable that they thrived at depths of 25 cm, where they can out-compete other phototrophs. *Prosthecochloris* have been previously observed in many marine and saline habitats, such as the Black Sea [59], Baltic Sea, Sippewissett Salt Marsh, and Badwater basin [52]. They are considered to belong to a specialized phylogenetic lineage of green sulfur bacteria adapted for marine and salt water ecosystems. Blooms of *P. vibrioformis* have been previously observed in stratified lakes, where they dominate the community at a specific depth [60], sometimes forming clonal blooms [61].

The suspended phototrophic bloom was spatially organized analogous to benthic phototrophic mats in the nearby Sippewissett Salt Marsh [62–64] and elsewhere [65, 66]. Thus, the disturbance experiment that we performed *in situ* created a transient ecosystem with niches resembling those in coastal microbial mats, except across a spatial scale that was one order of magnitude greater. The community slowly collapsed after about two weeks and the water column seemed to return to its original state (Figure 5). We did not observe a shift from phototrophic to chemotrophic sulfur oxidation after the phototrophic bloom [21].

**Figure 5:**
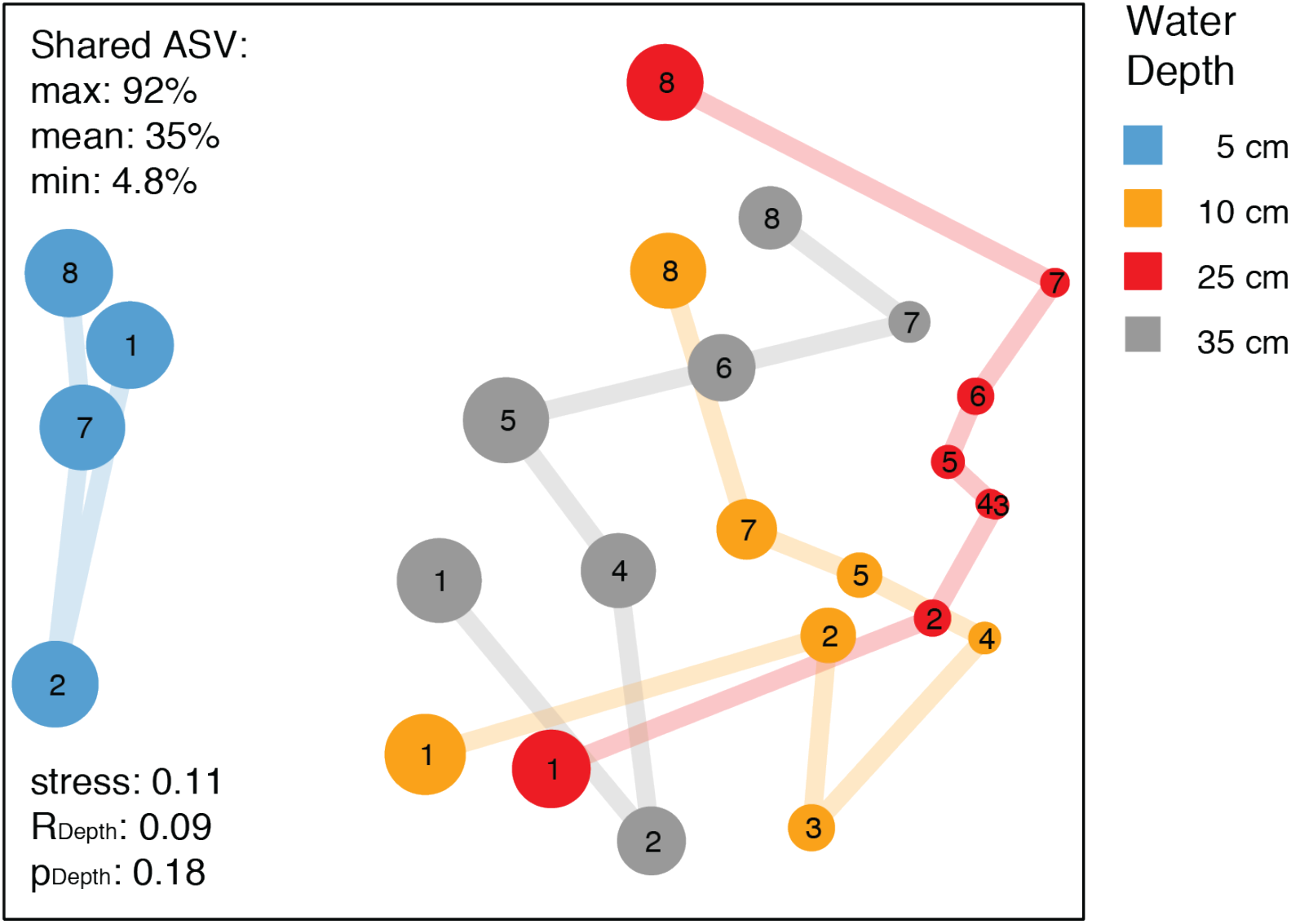
Microbial Community Turnover. Non-metric multidimensional scaling (NMDS) ordination based on relative abundance of ASV (amplicon sequence variants). Each circle represents one sample, the closer two samples are the more similar is their microbial community structure. Circle size represents Shannon Diversity. Numbers indicate sampling time points. Colors indicate bloom layers. Note: Individual holes were very similar (see Figure S8A) and thus we averaged relative ASV abundances for clarity, i.e. each circle represents an average across replicate experiments. NMDS ordinations for individual experiments are shown in Figure S7. The communities in the different layers of individual experiments are significantly different yet overlapping (see Figure S7).

### New species of green sulfur bacteria and possible viral predation

Due to the findings of a previous study based on 16S rRNA gene libraries, Imhoff and colleagues proposed the existence of several uncultivated GSB species in Sippewissett Salt Marsh and other estuaries [52]. The authors provide evidence that several GSB clades harbor species that have yet escaped isolation, among those are species in the genera *Chlorobaculum* and *Prosthecochloris*.

We have strong evidence that we found at least two of these species based on our MAGs of a *Chlorobaculum* species (Bin 6, Figure S13, S15) and a *Prosthecochloris* species (Bin 10, Figure S13, S16). Both MAGs cluster sufficiently far away from the closest cultured isolate (Figure S12, S14) and have average nucleotide identity (ANI) values of <90 to their respective closest cultured isolate.

Physiological and functional characteristics cannot be easily deduced from the presence of genes alone. Yet, it is very likely that the populations represented by bin 6 and 10 were sulfur oxidizers, because their genomes encoded for enzymes performing thiosulfate oxidation. As in most GSB, bin 6 contained the genes SoxABXYZ coding for enzymes that oxidize thiosulfate to sulfate and polysulfides [67]. Bin 10 only contained SoxYZ (Figure S17). The absence of SoxB genes has been identified in other non-thiosulfate oxidizing GSB such as the close relative *Prosthecochloris estuarii*, or in *Chlorobium limicola* DSM 245 and *Chlorobium luteolum* DSM 273 [68]. Bacteriochlorophyll synthesis genes were identical in both Chlorobi MAGs coding for pigments common to *Chlorobi*. In bin 6 we found complete operons encoding for cytochrome o oxidase (CyoABCDE) and cytochrome d oxidase (CydAB) [69]. The latter was found also in bin 10, indicating that both organisms have means to cope with oxygen stress, hence explaining their presence at relatively high oxygen concentrations and their ability of performing anoxygenic photosynthesis under (hyp)oxic conditions.

Both MAGs also contain CRISPR-Cas systems that are very different from the cultured isolates (Figure S18, S19). Our CRISPR results show that Trunk River populations may be under predatory stress, affecting the abundance of bacterial blooms, and that host immunity is active in this ecosystem [70]. The unique CRISPR arrays indicate that closely related species may be infected by different viruses with species specificity [71]. However, some viral populations have been reported to have broad host ranges [72]. Divergent evolution or strain level microdiversity may also explain distinct CRISPR-Cas systems [73]. A lack of public databases containing viral sequences restricts the detection of viral-host interactions [74]. While Llorens-Marès *et al.* (2017) characterized a potential green sulfur bacterial viral infection, to date, phages infecting *Chlorobi* have not been reported. Our analyses suggest that viruses of the family *Microviridae* played a major role in the transient bloom (Figure S19), possibly even for its demise. This interesting finding merits future research on transient phototrophic blooms in estuarine ecosystems.

## Conclusions

In this study, we investigated phototrophic blooms that naturally occur in a brackish estuarine ecosystem to understand the underlying microbial and biogeochemical dynamics. We show that phototrophs belonging to different phyla spatially organized within the water column based upon their light and oxygen requirements forming a layered bloom, analogous to the layered communities in phototrophic microbial mats (Figure 7). The community composition of the bloom indicates the presence of a sulfur cycle between anoxygenic phototrophs and sulfur reducers that could explain the rapid development of the bloom. We identified metagenome assembled genomes of two novel species of green sulfur bacteria, belonging to *Chlorobaculum* and *Prosthecochloris*. The MAGs contained terminal oxidases that may allow the organisms to thrive in a hypoxic environment, as well as repeats that evidence of host-virus dynamics. The metagenome contained viral sequences suggesting that *Microviridae* viruses infect species within the *Chlorobiales*. Overall, Trunk River is an excellent model ecosystem to study the microbial ecology and ecophysiology of phototrophic blooms in a natural setting.

**Figure 6:**
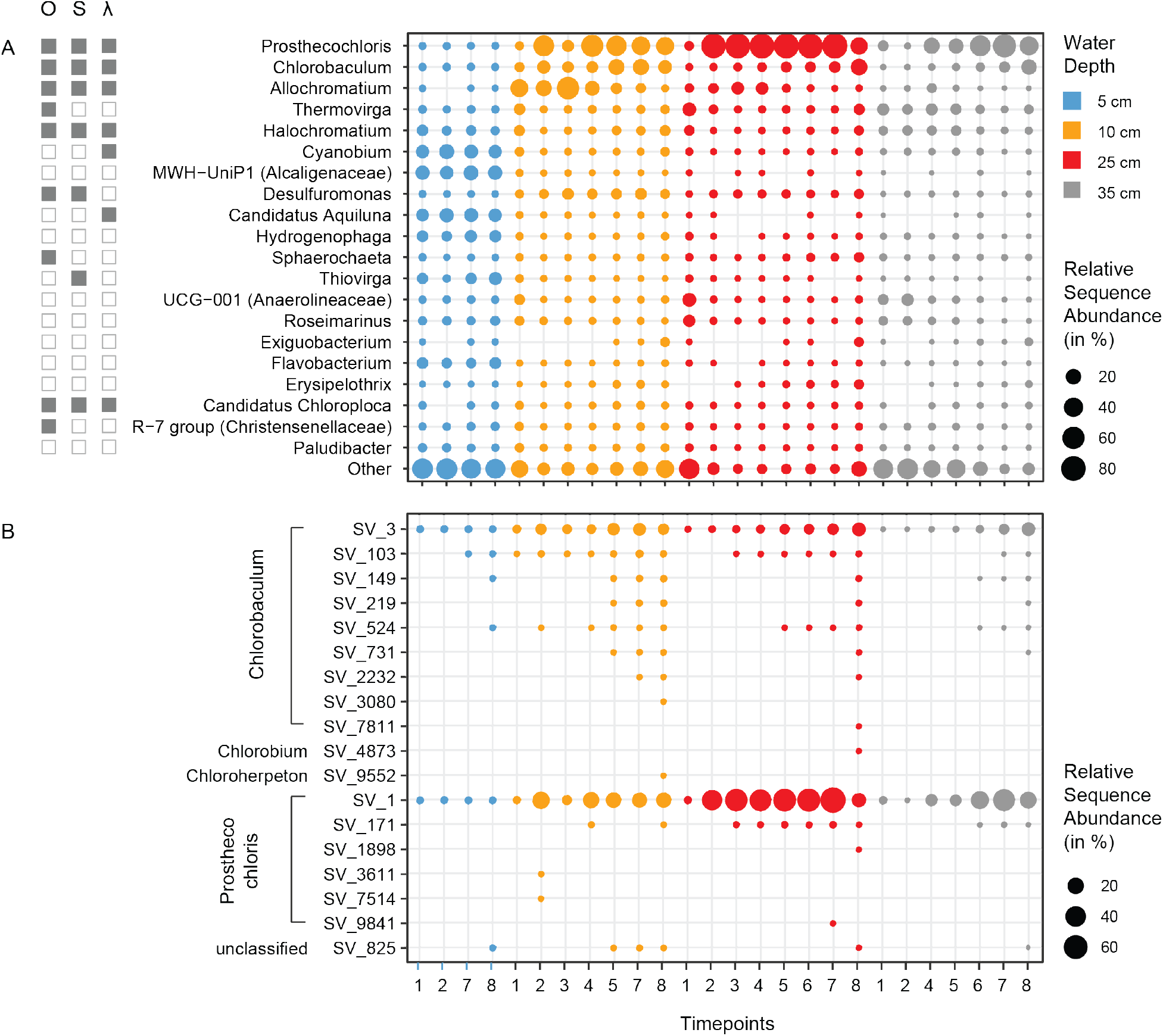
Microbial community composition on genus level. A: Relative sequence abundance of genera found in different depths (colors) and timepoints (x-axis). Relative sequence abundances were averaged across triplicates, due to the high similarity of all three experiments. Clades that are anaerobic (O), involved in the sulfur cycle (S), or phototrophic (ƛ) are indicated by full squares. B: Relative sequence abundance of amplicon sequence variants (ASV) within the order *Chlorobiales*. The graph shows average values of the three replicate experiments for clarity. The replicate experiments were very similar (see SI Figure S5 and S6).

**Figure 7:**
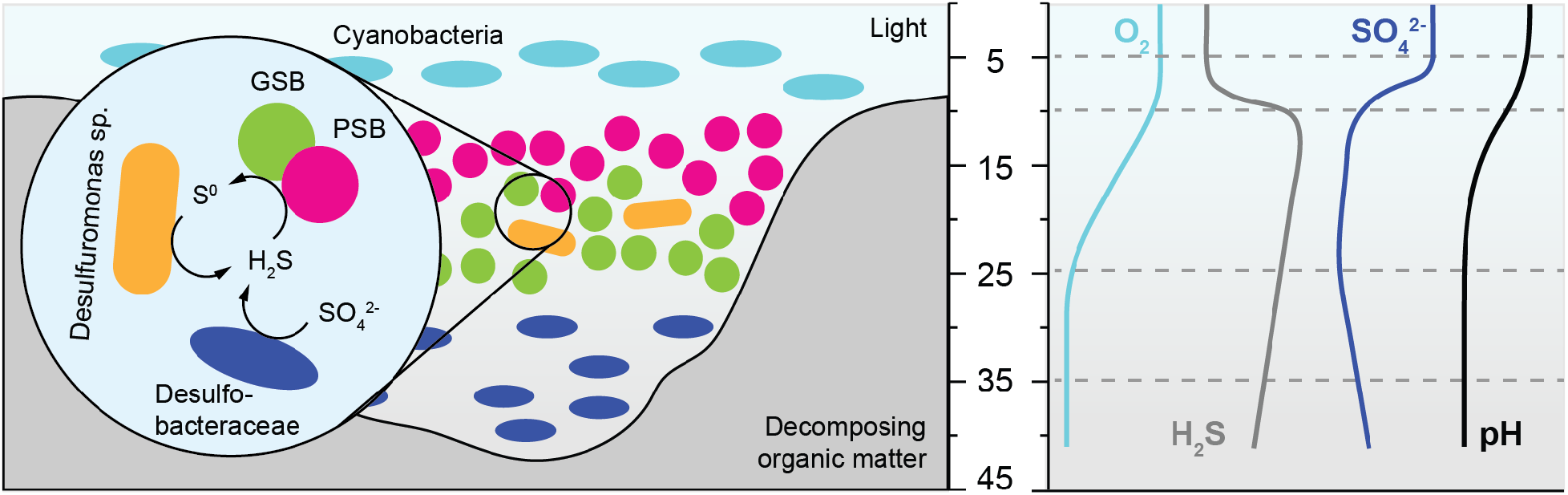
Schematic overview of the phototrophic bloom showing the main phototrophic populations, sulfur compounds, and chemical gradients.

## Methods

### Experimental Setup and Sample Collection

We used custom-made sampling poles for long-term environmental monitoring of the water column without disturbing the established gradients (LEMON; Figure 1B, C). The sampling poles were placed in three replicate depressions (A-hole, E-hole, and K-hole) that we dug into the thick layers of decaying organic matter (Figure 1A). In each of sites, a sampling pole was placed such that the inlets sampled water at 5 cm, 10 cm, 25 cm, and 35 cm depth below the surface (Figure 1B, C). Sampling poles were set up one day after the holes were dug out and sampling began one day after set up (two days post disturbance), to allow disturbed sediment to settle. Samples were collected over a 15-day period during July-August 2015. For each sample, the first 5 ml were discarded, followed by collection of 100 ml of water in several sterile tubes for further analyses. The tubes were transported on ice to the laboratory and stored at 4°C. All sample collections were carried out between 4 pm and 6 pm.

### Enrichment cultures

To enrich for GSB we used a defined saltwater medium (400g/l NaCl, 60g/l MgCl_2_*6H_2_O, 3g/l CaC_l2_*2H_2_O, 10g/l KCl) buffered at pH 7.2 with 5 mM MOPS. The medium contained 5 mM NH_4_Cl as N source, 1mM K phosphate (pH 7.2) as P source, 70 mM NaHCO_3_ as C source, 10 mM Na_2_S_2_O_3_ as electron donor, 1 mM Na_2_S as reductant or electron donor, a multivitamin solution prepared at 1000× in 10 mM MOPS at pH 7.2, and a trace metal solution prepared at 1000× in 20 mM HCl. Saltwater base, MOPS, N- and P-source, and trace metals were autoclaved together in a Widdel sparging flask, cooled under a stream of N_2_/CO_2_ (80%:20%) gas. C-source, electron donors and vitamins were added from filter-sterilized stock solutions after cooling. The medium was inoculated with biomass removed from *in situ* enrichments of GSB grown on glass slides using a 770 nm monochromatic LED. After inoculation, the bottle was kept in dark for 2-4 hours and then placed 5 cm away from a LED light source with the same specifications. After a visible sign of growth – green coloration – the culture was filtered through 0.2 µm filter and used for DNA extraction, similar to other samples.

### Physicochemical Measurements

*In-situ* measurements of pH, temperature, dissolved oxygen, oxidation reduction potential (ORP), and ion selective electrode (ISE) were carried out with a multi-parameter probe equipped with a quarto probe (YSI Professional Series Model Pro). The probe was calibrated for pH with 4, 7, and 10 buffers and for dissolved oxygen using oxygen-saturated water and an anoxic solution of sodium ascorbate and sodium hydroxide. After each sample collection the probe was lowered into the water to each depth per site and after probe readings stabilized, the parameters were recorded.

To measure biomass and pigment spectra, up to 10 ml of the collected sample was filtered through a sterile Millipore filter (0.2 µm GTTP, 0.2 µm GNWP, or 0.22 µm GV). Filtrates were washed twice with ammonium acetate solutions with the same ionic strength as each depth. The filters were placed on aluminium foil, dried at 60°C overnight and subsequently weighed (Figure S3). A Spectral Evolution SR1900 spectrophotometer was used to measure the spectrum of the dried biomass on each filter with a scanning range of 350-1900 nm. The light source was a Dyonics 60 W lamp.

After sterile filtration, the filtrate was used to measure anion, cation, and organic acid concentrations using an ion chromatographer. The ion concentrations of samples were measured by diluting filtrate 1:10 with Millipore water to a total volume of 2 ml. The diluted samples were measured in triplicate using a ThermoFisher/Dionex ICS2100 equipped with an AS18 column using a 13 minute, 33 mM NaOH isocratic program to measure anions and a CS12A column using a 13 minute, 25 mM methane sulfonic acid isocratic program to measure cations. Samples for organic acid analysis were filtered through 0.2 μm filters and 900 μL of filtrate was added to 100 uL of 5 M H_2_SO4 to precipitate any compounds that might do so on the column. The samples were centrifuged and the upper portion was removed for HPLC analysis. Samples were analyzed on a BioRad Aminex HPX-87H column in isocratic elution mode with 5 mM sulfuric acid.

Iron concentration was quantified using the ferrozine assay [75]. 4.5 ml filtrate were added on site to 0.5 ml of 1 M HCl to prevent oxidation of any available Fe(III). For Fe(II), 50 µl filtrate was added to 50 µl of 1M HCl and 100 µl of ferrozine (0.1% [wt/vol] in 50% ammonium acetate) was added. For total iron, 50 µl filtrate was added to 50 µl of 10% hydroxylamine hydrochloride in 1M HCl to reduce Fe(III) to Fe(II). Samples were added to 100 µl of ferrozine. All samples were incubated for 15 min and a filtrates Absorbances were read in triplicate at 560 nm using a Promega plate reader. Ferrous ammonium sulfate was used as standard.

Sulfide concentrations were quantified using the Cline assay [76]. 1.5 ml filtrate were added on site to 500 µl of zinc acetate solution (91 mM) to prevent oxidation of the sulfide. Cline reagent (N, N-dimethyl-p-phenylenediamine sulfate, H_2_SO_4_, NH_4_Fe(SO_4_)_2_·12 H2O) was added, the samples were incubated in the dark for 30 minutes and absorbance was read at 665 nm. A table with all physicochemical and biomass measurements is publicly available at PANGAEA (https://doi.pangaea.de/10.1594/PANGAEA.900343)

### DNA Extraction, Library Preparations, and Sequencing

Within 2 – 6 hours of sample collection, 50 ml sample was filtered using an autoclaved 0.2 µm polycarbonate filter (GTTP Millipore) and stored at −20°C. Each filter was cut with a sterile blade and extracted with the MoBio PowerFecal kit. We followed the protocol, but instead of bead beating, the samples were twice vortexed horizontally with the beads (10 min and 20 min with a 10 min pause). DNA concentration and purity were measured with Nanodrop and Promega Qubit fluorometer and Nanodrop, respectively.

We prepared 16S rRNA gene amplicon library using V4-V5 fusion primers as previously described [77]. Briefly, the fusion primer contains TruSeq adapter sequences, barcodes, and the forward or reverse 16S rRNA gene primers. The forward and reverse 16S rRNA gene primers were 518F (CCAGCAGCYGCGGTAAN) and 926R (CCGTCAATTCNTTTRAGT). The PCR conditions were as follows: initial denaturation of 94°C for 3 min, 30 cycles of denaturation at 94°C for 30 s, annealing at 57°C for 45 s, extension at 72°C for 1 min, and final extension at 72°C for 2 min. The libraries were cleaned using Agencourt Ampure XP beads, quantified using picogreen, pooled in equimolar ratios, and cleaned again using Agencourt Ampure XP beads a second time. The indexed libraries were then sequenced on the Illumina MiSeq PE250 platform.

The DNA from the depth of 25 cm from timepoint 7 from the three replicates, as well as from a phototrophic enrichment were used to generate whole-genome shotgun metagenomic library. The DNA was sheared using Covaris sonicator, size selected for 500-600bp using Pippin prep, and cleaned using Agencourt Ampure XP clean beads. The cleaned DNA was analyzed using Bioanalyzer DNA1000 chip and used to prepare metagenomic library using Nugen Ovation ultralow DR multiplex kit with manufacture supplied protocol. The libraries were then sequenced on Illumina MiSeq PE250 platform. All the sequencing was performed at the Keck facility at the J. Bay Paul Center at the Marine Biological Laboratory, Woods Hole, MA.

### Amplicon Sequence Data Analyses

The amplicon data was demultiplexed in mothur v1.39.5 [78], followed by the trimming of 16S rRNA gene amplification primers using Cutadapt v1.16 [79] with default parameters. The primer-trimmed amplicon sequencing data was quality checked using DADA2 v1.9.0 R Package [80]. In DADA2, the reads were trimmed at the first instance of quality drop below 8, an expected error rate of 2, followed by trimming to 220bp and 200bp for forward and reverse reads. Any reads that matched PhiX or had an ambiguous base were removed. An error profile for the forward and reverse reads was generated using learnErrors function and then used to merge the forward and reverse reads using the mergePairs function. The merged reads were used to generate the amplicon sequence variants using makeSequenceTable function, which was then filtered for chimeras using removeBimeraDenovo function. The amplicon sequence variants were assigned taxonomy in DADA2 using Silva reference database v132 [81]. Community analyses were performed using a custom workflow based on R and the packages vegan, labdsv, tidyverse (stringr, dplyr, ggplot2), UpSetR and custom scripts [82–87] for details see. Relative abundance of bacterial ASV (amplicon sequence variants), Bray-Curtis dissimilarities, Nonmetric Multidimensional Scaling as well as analyses determining Singletons and percent shared ASVs are based on the unaltered Sample×ASV table as calculated by DADA2. The ASV×Sample table including taxonomy is available at PANGAEA (https://doi.pangaea.de/10.1594/PANGAEA.900354). To compare the diversity between samples using the number of observed species, Shannon index, Inverse Simpson diversity and Chao1 Richness [88] the ASV abundance tables were rarefied to account for unequal sampling effort using 31,682 randomly chosen sequences without replacement. For details refer to the R workflow available at the public database PANGAEA (https://doi.pangaea.de/10.1594/PANGAEA.900344).

### Metagenomic Sequence Data Analyses

Quality control of the raw reads was performed using Preprocessing and Information of SEQuence data (PRINSEQ) to remove sequencing tags and sequences with mean quality score lower than 25, duplicates and N’s [89]. All runs combined provided a total of approximately 3.5 million 250 bp read pairs. All forward and reverse reads were placed together in one file and cross co-assembled using SPAdes using the --meta option [90]. Binning was performed using MetaBAT [91] and Anvi’o (v5.2) metagenomic workflow (CONCOCT) [92]. Completeness and contamination of bins was assessed using CheckM [93]. Assembled genomes that contained more than 90% genome completeness, less than 5% contamination, and sequences mainly from a single genus were further analyzed. This yielded two high quality bacterial metagenome-assembled genomes (MAGs): Bin 6 and Bin 10. Taxonomic composition for each bin was predicted using FOCUS [94]. Phylogenetic analysis including the identification of their closest phylogenetic neighbors was investigated using PATRIC Comprehensive Genome Analysis [95]. Gene prediction for MAGs was performed using prodigal (V2.60, -p meta). We searched for sulfur, terminal oxidases and chlorophyll pathways using Ghost-KOALA against the KEGG GENES database. The *Chlorobi* bins 6 and 10 contained 2008 and 1938 predicted proteins, respectively. CRISPRCasFinder [96] and CRISPRone [97] were used to identify CRISPR repeat and spacer sequences. The quality checked reads from each sample were mapped to the MAGs, Bin 6 and Bin 10 using bowtie2 [98]. The mapped reads were then analyzed using iRep [42] to estimate replication events in Bin 6 and Bin 10. Unassembled sequences were processed on the MG-RAST platform version 4.0.3. Percent abundance of viral sequences was calculated from the RefSeq database using an e-value cutoff of 1e-5, a minimum identity cutoff of 60%, and an alignment length minimum cutoff of 15 [99]. For details refer to the metagenome analyses workflow publicly accessible at HackMD (https://hackmd.io/tGZyCM9sSNmuorpHenQVNA).

## Declarations

### Availability of data and material

The genomic datasets generated and analyzed during the current study are available on MG-RAST (Project Name: Trunk River, ID: 4837589.3 (sample SK), 4837590.3 (sample 7K3), 4837591.3 (sample 7E3), 4837592.3 (sample 7A3)) and metagenome-assembled genomes workflow are available on HackMD (https://hackmd.io/tGZyCM9sSNmuorpHenQVNA). The raw 16S rRNA amplicon data, the shotgun metagenomic data, the 16S rRNA gene clonal sequences, and the metagenome assembled genomes presented in this work are publicly archived in NCBI under Bioproject PRJNA530984 (https://www.ncbi.nlm.nih.gov/bioproject/530984). The contextual datasets generated and analyzed during the current study are publicly available at PANGAEA under: https://doi.org/10.1594/PANGAEA.900343

### Authors’ contributions

SB helped design the study, collected samples, prepared and analyzed sequencing data, wrote manuscript

ESC helped design the study, collected samples, prepared and analyzed physicochemical data, wrote manuscript

SHK helped design the study and enrichment cultures, analyzed physicochemical data and pigment spectra, wrote manuscript

SPC analyzed metagenomic data, wrote manuscript

SK obtained enrichment culture, wrote manuscript

SD helped design the study, wrote manuscript

KH collected samples, prepared and analyzed physicochemical data and pigment spectra, wrote manuscript

SER designed the study, analyzed and visualized sequencing and physicochemical data, wrote manuscript with input from all co-authors

All authors read and approved the final manuscript.

## Acknowledgements

We would like to express our deepest gratitude towards Dianne Newman and Jared R. Leadbetter, directors of the Microbial Diversity Summer Course at the Marine Biological Laboratory 2014-2017. Without their trust, support and encouragement the project would not have been realized. We are very grateful to Joseph Vinneis, Kim Finnegan, and Mitchell Sogin for providing laboratory space, equipment, and guidance regarding next-generation sequencing. We are also very grateful to the students and staff of the 2015 Microbial Diversity Summer Course, specifically Kurt Dahlstrom, Kristina Garcia, Jessica Choi, Rachel Soble, and Lina Bird. We would like to thank the Simons Foundation, NSF, DOE, and NASA for funding the Microbial Diversity Summer course, as well as Promega, Thermo Fisher Scientific, Spectral Evolution, and MBL’s Marine Resources Center for providing reagents and equipment used in this work.

