## Supplementary Information for "Microbial community dynamics and coexistence in a sulfide-driven phototrophic bloom"

Supplementary Materials and Methods

<u>16S rRNA gene libraries and construction of phylogeny</u>

From the DNA of Sample 7A3 (Replicate hole A, at 25 cm depth at time point 7) clone library was made in following way. The 16S rRNA gene was amplified using 27F (Bacterial - AGAGTTTGATCMTGGCTCAG) or 4Fa (Archaeal - TCCGGTTGATCCTGCCRG) forward primer and 1391R (broad range - GACGGGCGGTGTGTRCA) reverse primer. The PCR product was cleaned using Promega Wizard® PCR Cleanup system and cloned using pGEM®-T easy system using manufacturer supplied standard protocol (Promega Corporation, Madison, WI). Briefly, the cleaned PCR product was ligated in vector, followed by vector cloning into Promega JM109 competant cells. The transformed cells were selected and screened using blue-white colony screening on LB plate with ampicillin, X-gal, and IPTG. The transformed cells were grown in liquid LB broth with ampicillin for 12 hours and the plasmid was extracted using Promega PureYieldTM plasmid system. The plasmid was send to Sequetech DNA Sequencing Service (Mountain View, CA) for Sanger sequencing from one end using the T7 primer.

16S rRNA gene phylogeny was calculated using ARB/SILVA (Ludwig et al., 2004) and the SILVA database SSU Ref NR 132 (Quast et al., 2013). We added the sequences to the SILVA tree using the ARB maximum parsimony quick-add tool and chose near full-length (>1300 nucleotides) reference sequences from neighboring clades and isolates. We aligned these sequences using SINA (Pruesse et al., 2012) and manually curated the alignment based on ribosomal secondary structure. This alignment was used to calculate a tree with the RAxML8 algorithm (Stamatakis, 2014) and a positional variability filter including only conserved positions in the alignment with ≤3.1% mutation rate. The final alignment had 1007 valid columns encompassing the E. coli reference positions 1785-40339. The tree was calculated with 100 iterations of which the most robust tree was selected. The partial gene sequences from this study were added to the consensus tree using the same positional variability filter without changing the overall tree topology.

### Supplementary Results

Visual Observations

At the first sampling time point (2 days post-disturbance), no difference was observed in the water column or in the samples collected from different depths. Two days later (time point 2, 4 days post-disturbance), a faint pink layer was observed in the water column, and faint shades of yellow in samples from the 25 cm depth (Figure S2). At the next sampling time point (time point 3; 5 days post-disturbance), we observed a bright yellow suspension below the water surface (Figure 1). Of the samples collected at this time point, the samples from 25 cm depth were most distinctly yellow (Figure S2). The yellow color of the suspension intensified by time point 4 (7 days post-disturbance). No remarkable visual changes in the system were observed for the subsequent three time points (time points 5, 6, and 7). During these three time points, the yellow suspension only slightly changed. At time point 8 (16 days post-disturbance), the holes had partially collapsed, the water was much clearer and the 25-cm samples showed a reduction in the intensity of the yellow color (Figure S2). We found a green layer at the bottom of each hole, seemingly from sedimented GSB. It has to be noted that the opacity of the samples was higher at the first two timepoints than at the following timepoints. Especially at timepoint 1 the suspension was beige and very opaque, while later on the suspension became more yellow, but also more translucent (Figure S2).

Physicochemistry

Within the first three days the pH decreased between 1-2 units in all layers, from around pH 9 to pH 7 at 5 cm water depth (7.7±1.0), from pH 8 to pH 7 at 10 cm (7.3±0.7) and from pH 8 to pH 6.3 at 25 (6.7±0.8) and 35 cm (6.8±0.7) (Figure 2). Over the 15-day sampling period, the pH at 5 cm (7.3±0.8) and 10 cm (6.9±0.6) water depth showed more variation than at 25 cm (6.5±0.6) and 35 cm (6.5±0.5). At depths within and below the bloom, pH was very constant between 6 and 6.3.

The Fe(II) concentration at 5 cm and 10 cm was < 5 µM (Figure S4). The Fe(II) concentration at these depths showed less fluctuation over the 15 days of sampling as compared to 25 cm and 35 cm. With an average range of 10 – 12 µM concentration, samples from 25 cm had the highest concentration of Fe(II). When compared to Fe(II), the Fe(III) concentrations were consistently lower at all depths between 1-5 µM. With roughly 20-25 µM of total iron, 25 cm samples had the highest iron concentration (Figure S5).

Nitrate concentration showed more fluctuation over the sampling period at shallower depths, specifically at 5 cm and 10 cm, with a short-lived nitrate spike 5 days after the start of the experiment (Figure S5). This spike coincides with a concentration dip in the major ions suggesting an input of nitrate rich freshwater, potentially from fertilizer-rich soil runoff. The nitrate concentrations at 25 cm were slightly higher than at other depths in the water column. Ammonium and acetate concentrations were highest at 35 cm depth (up to 4 mM and 1.5 mM, respectively) with significantly lower concentrations at the shallower sampling points (25 cm, 10 cm, 5 cm) including no detectable ammonium throughout the entire experiment at the surface (5 cm). This indicates a source of ammonium and acetate at depth which is consistent with ammonification and acetate generation from fermentative organic matter degradation of the decaying seagrass underneath.

In addition, measurements of sodium, fluoride, chloride, bromide, phosphate, lithium, magnesium, lactate, formate, succinate, propionate, butyrate and glucose were made on all samples collected. The values are not shown for reasons of clarity, but the raw data are still available. The values for sodium and chloride were outside the range of the standard calibration curve, hence these values were not analyzed.

<u>Metabolic capabilities of Chlorobi MAGs (bin 6 and bin 10)</u>

Both MAGs contained genes of the thiosulfate-oxidizing multienzyme system (Sox). Bin 6 contained SoxXYZAB while Bin 10 contained only SoxYZ (Figure S15.5). Both MAGs also contained the sulfate reductase subunit DsrAB involved in sulfate reduction, and components of assimilatory sulfate reduction (CysND and Cys). Genes for bacteriochlorophhyll (BChl) biosynthesis (bchEMU) were found in both MAGs. Bd-type oxidases (CydAB) were present in both MAGS, while heme-copper oxygen reductases were only found in Bin 6 including seven cytochrome c oxidases (COX10, CyoABCDE and III) (Table S3.5).

The thiosulfate-oxidizing multienzyme (Sox) system is the widely distributed sulfur oxidation pathway found in SOB. Four protein complexes have been proposed in the Sox system: SoxAX, SoxB, SoxCD, and SoxYZ (Friedrich et al., 2001). SoxXA conjugates thiosulfate to the SoxY protein to form a cysteine S-thiosulfonate which is then degraded by SoxB (Sauvé et al., 2009). SoxCD genes oxidize thiosulfate to sulfate. The absence of SoxCD - as is the case for the here described *Chlorobi* MAGs - is characteristic for organisms that form sulfur globules (Welte et al., 2009). Substrates of the Sox system include thiosulfate, sulfide, elemental sulfur, sulfite, and tetrathionate.

Terminal oxidases have been suggested to play a role in oxygen defense in strictly anaerobic microorganisms (Ramel et al., 2013). They are divided into 1) heme-copper, 2) cytochrome bd, and 3) alternative oxidase families. The heme-copper family is divided into cytochrome c (cbb3) and quinol (Cyo) oxidases (Hamada et al., 2014). The bd-family of oxygen reductases has a wide phylogenetic distribution with homologs found in 18 bacterial phyla including *Chlorobi* and *Proteobacteria* (Borisov et al., 2011).

The chlorosomes (light harvesting complex) of GSB are divided into bacteriochlorophylls (BChls) *c*, *d* and *e*. GSB that are green in color produce chlorosomes containing either BChl *d* or *c*, and brown-colored GSB produce BChl *e* (Vogl et al., 2012).

Supplementary Figures

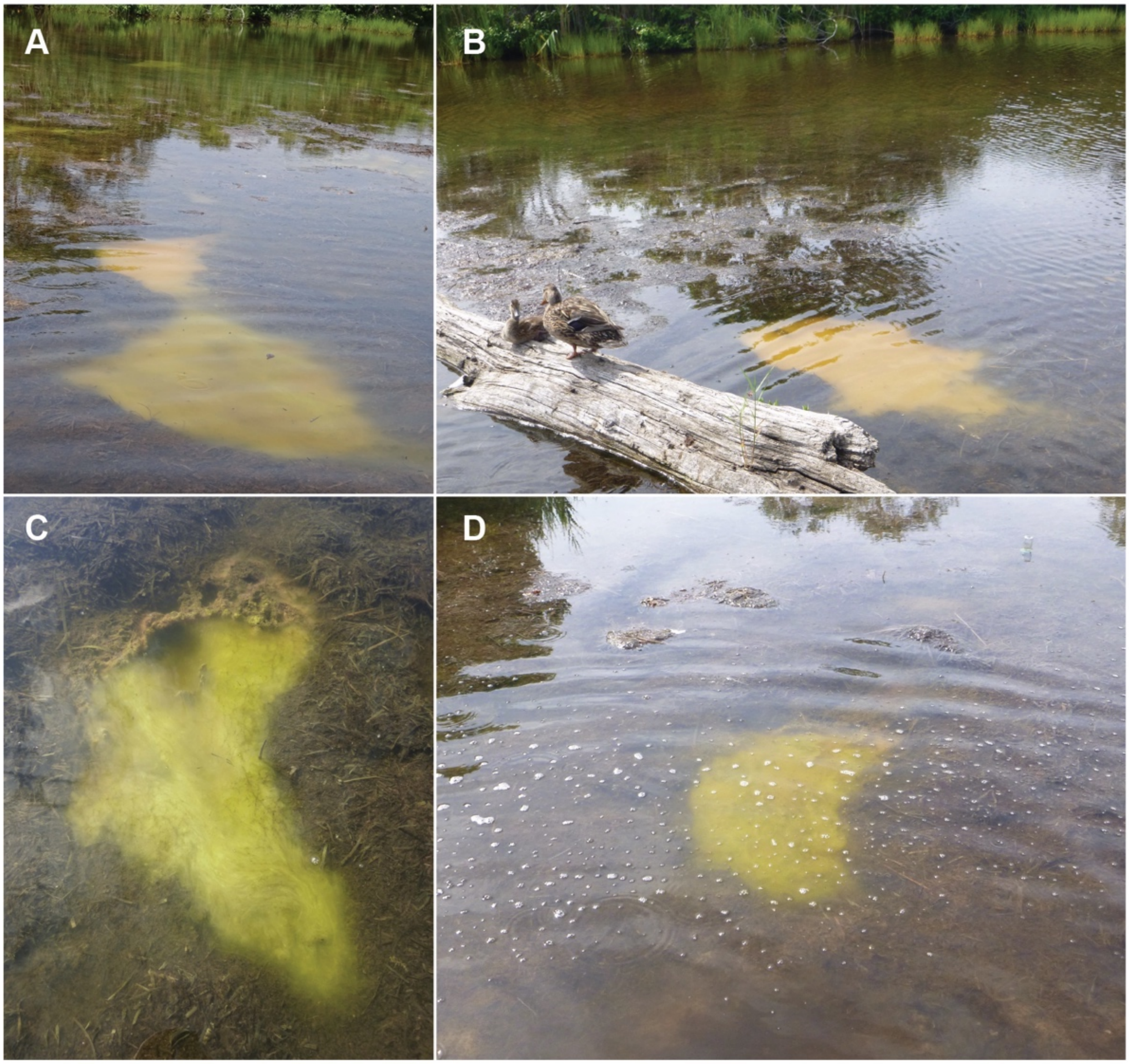

Figure S1: Pictures of natural blooms in Trunk River. B is the natural bloom shown in the aerial overview Figure 1A. C is a close-up of bloom D.

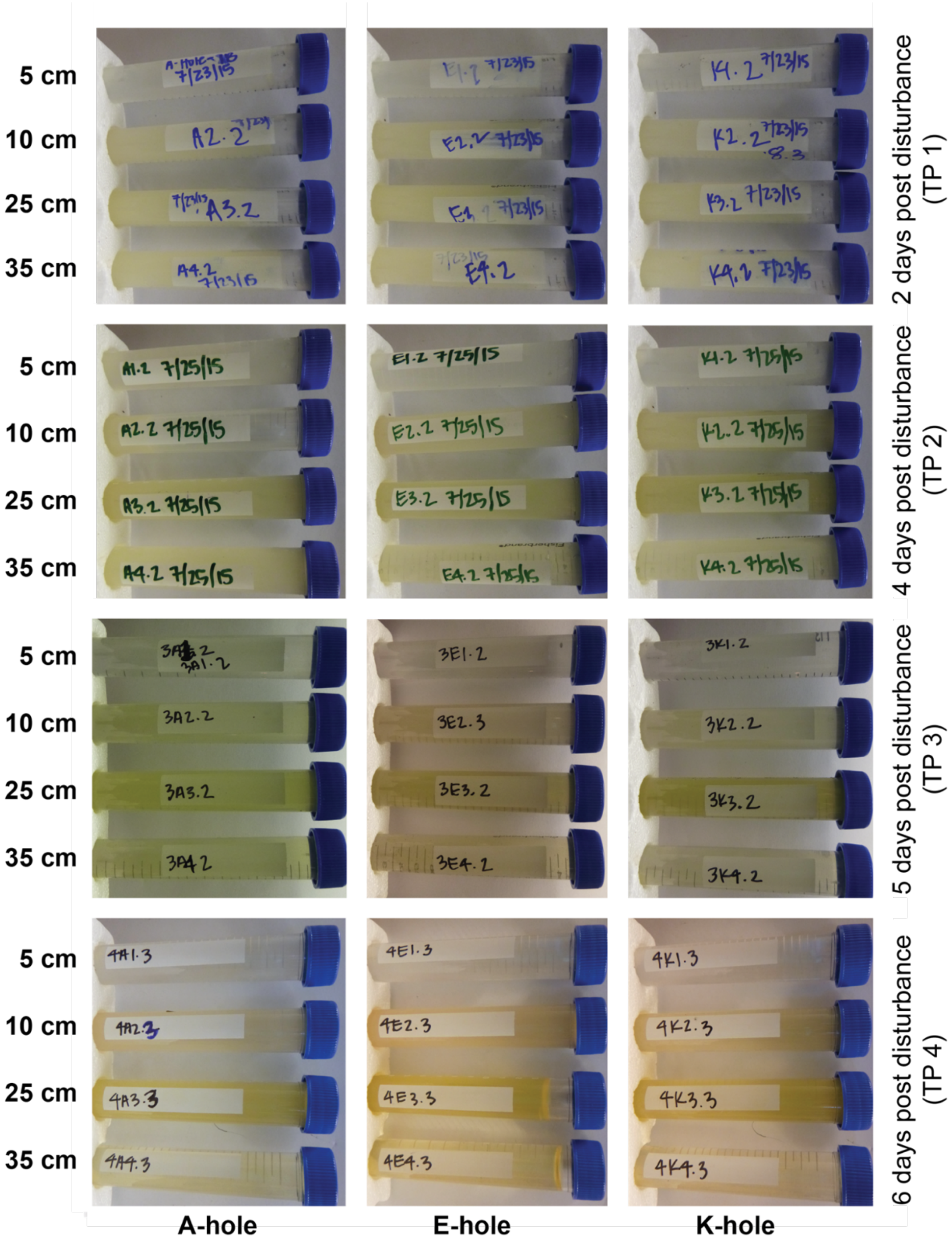
5 cm
10 cm
25 cm
35 cm
2 days post disturbance (TP 1)
4 days post disturbance (TP 2)
5 days post disturbance (TP 3)
6 days post disturbance (TP 4)
A-hole
E-hole
K-hole

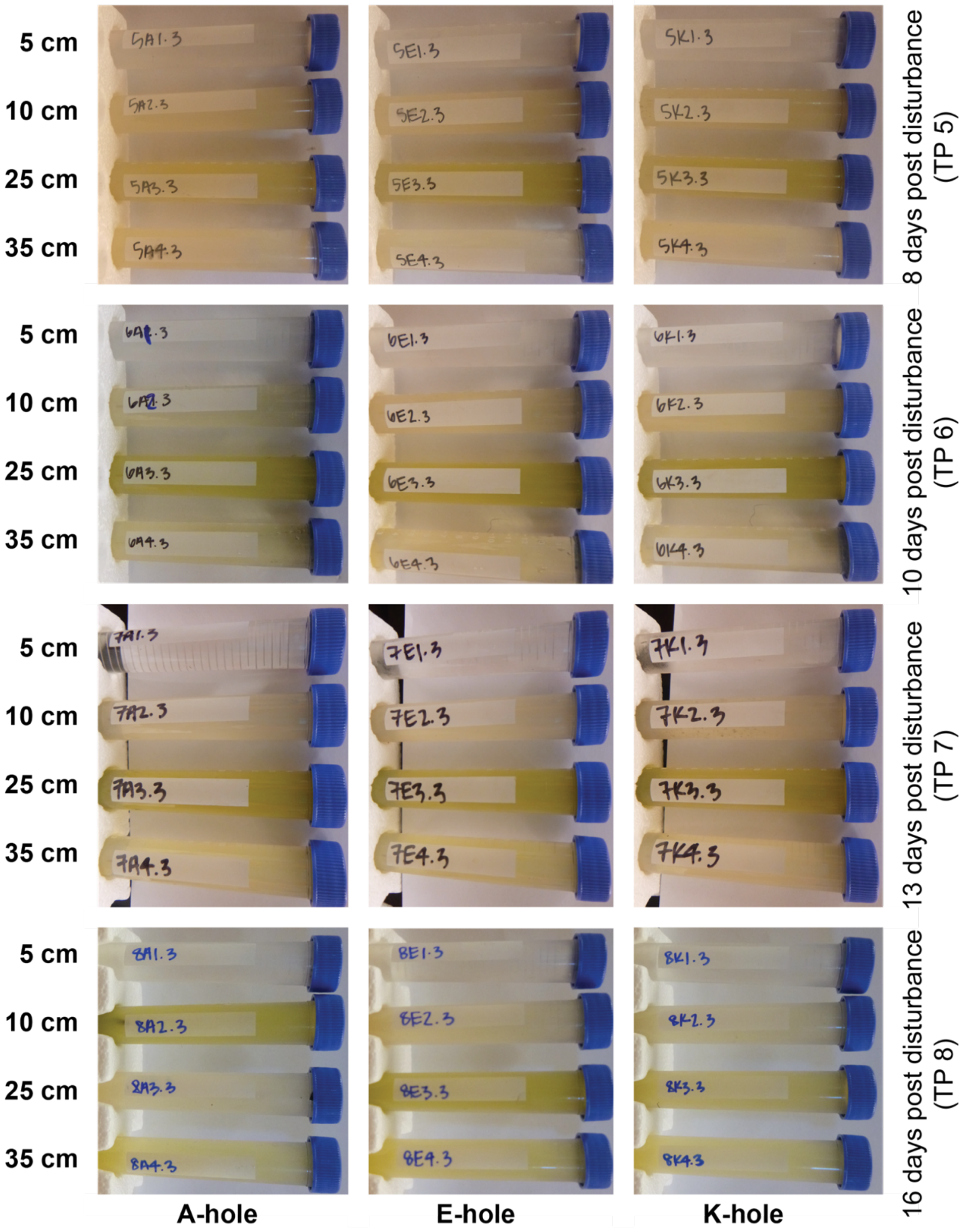

Figure S2: Color and appearance of samples from all holes, depths, and timepoints.

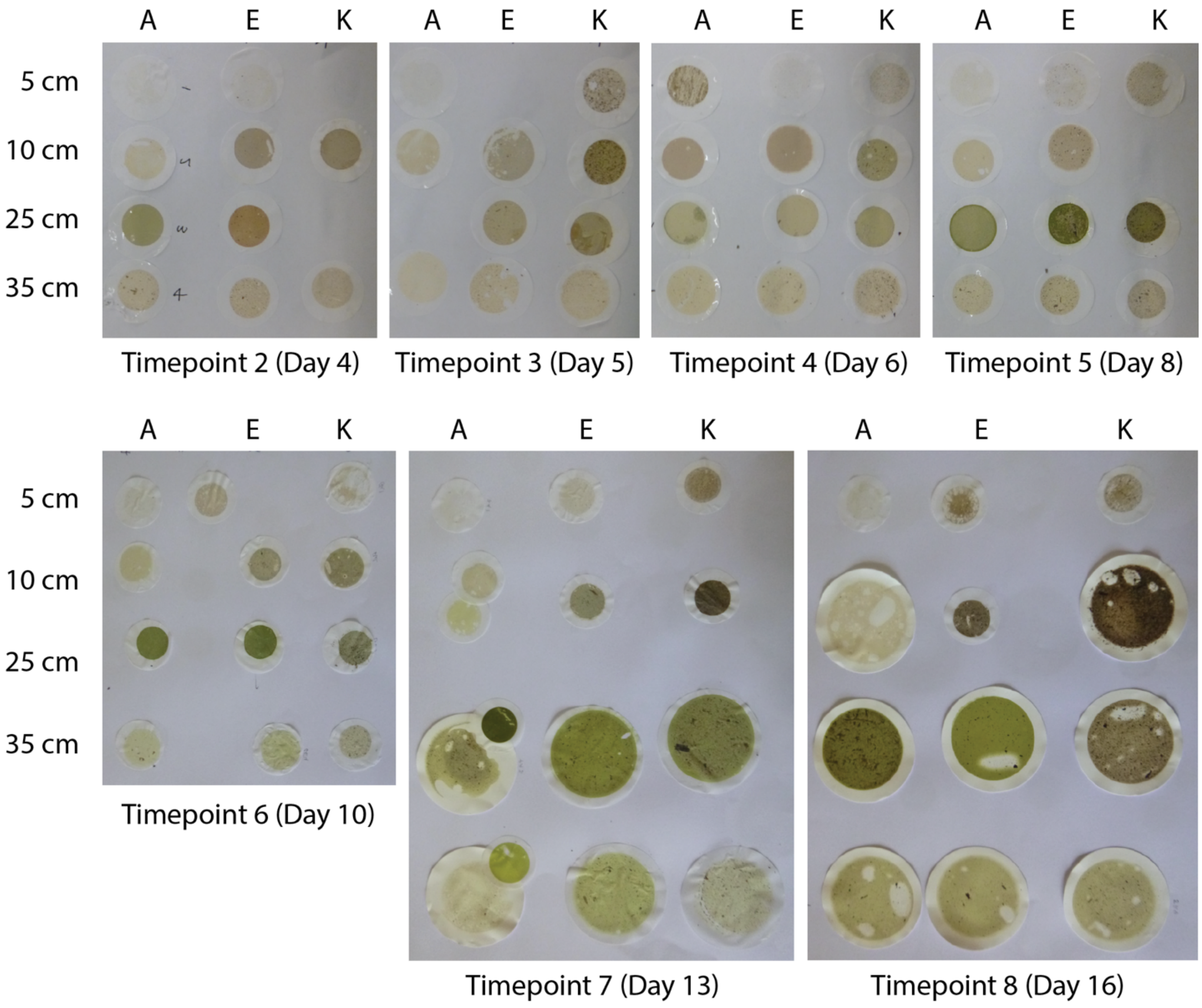

Figure S3: Filters that were used for biomass measurements and spectral analysis.

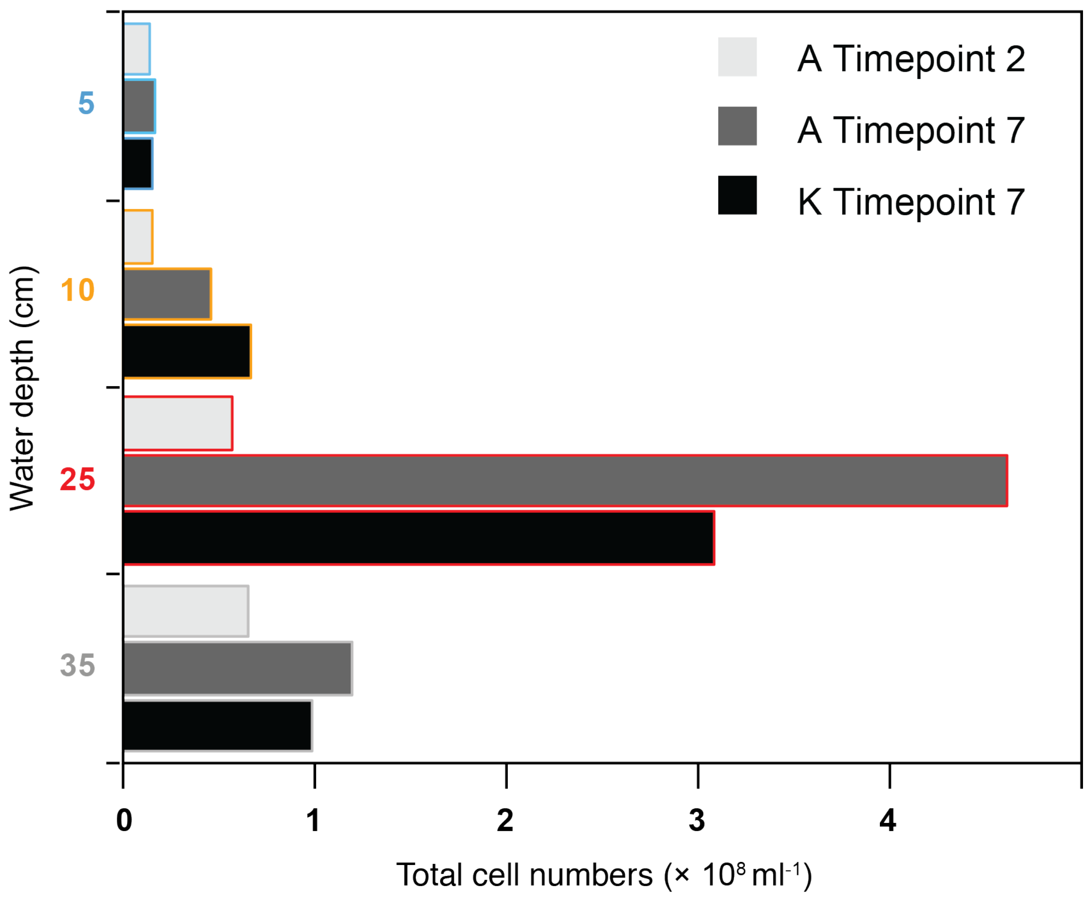

Figure S4: Total cell count of three samples (A2, A7 and K7) showing the differences in cell numbers between layers at the beginning and close to end of the experiment, corroborating the results obtained from biomass measurements.

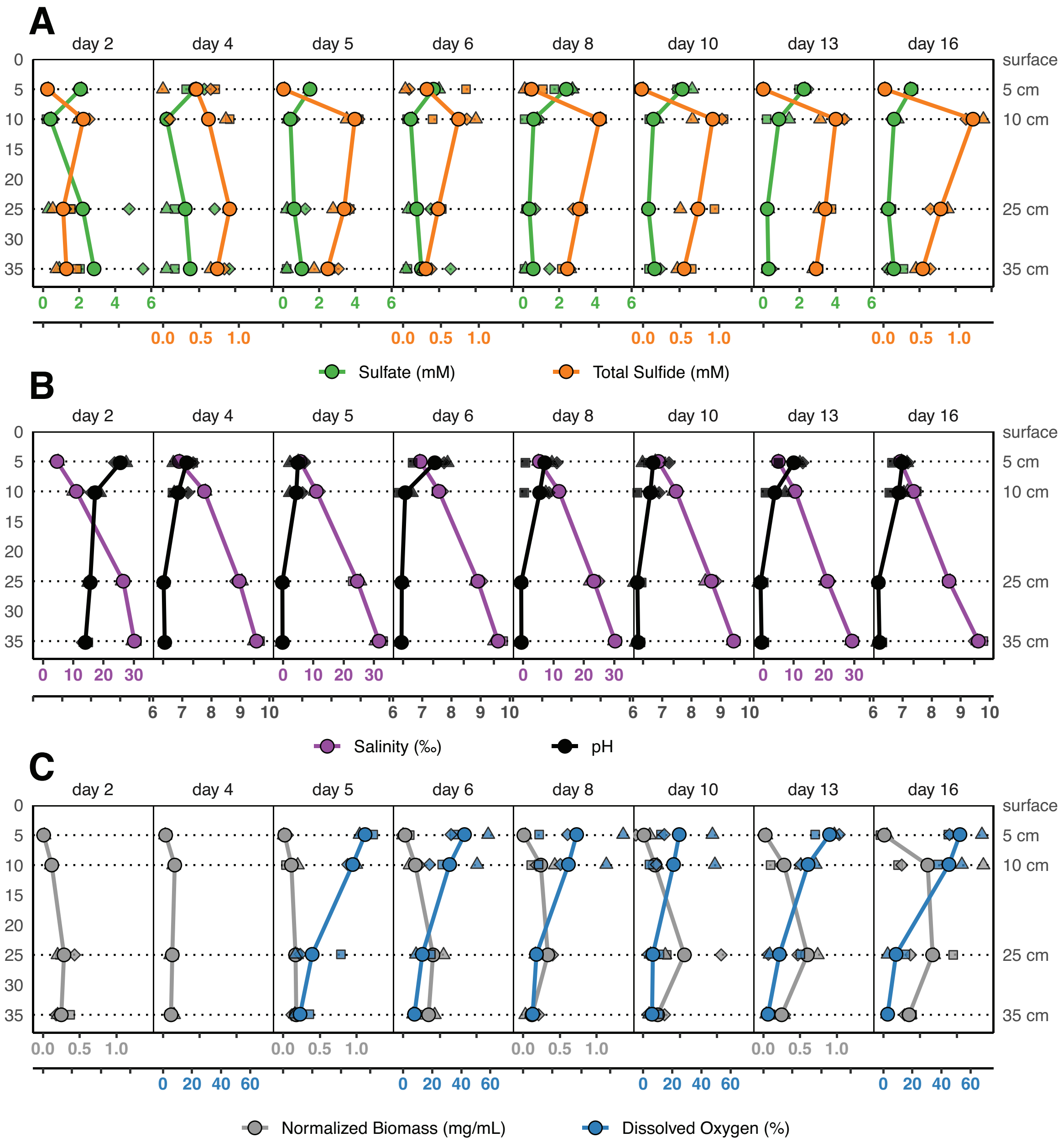

Figure S5: Depth profile representation of chemical data presented in Fig. 2. The main text figure (Fig. 2) was designed to focus on the best visualization of temporal trends in the data but makes it harder to intuit depth trends. This figure focuses on the best visualization of trends between the different experimental depths. A) sulfate and sulfide data. B) salinity and pH data. C) biomass and dissolved oxygen data. Circles and trend lines represent averages across all three experimental sites whereas smaller symbols (square, triangle, diamond) represent individual sample sites.

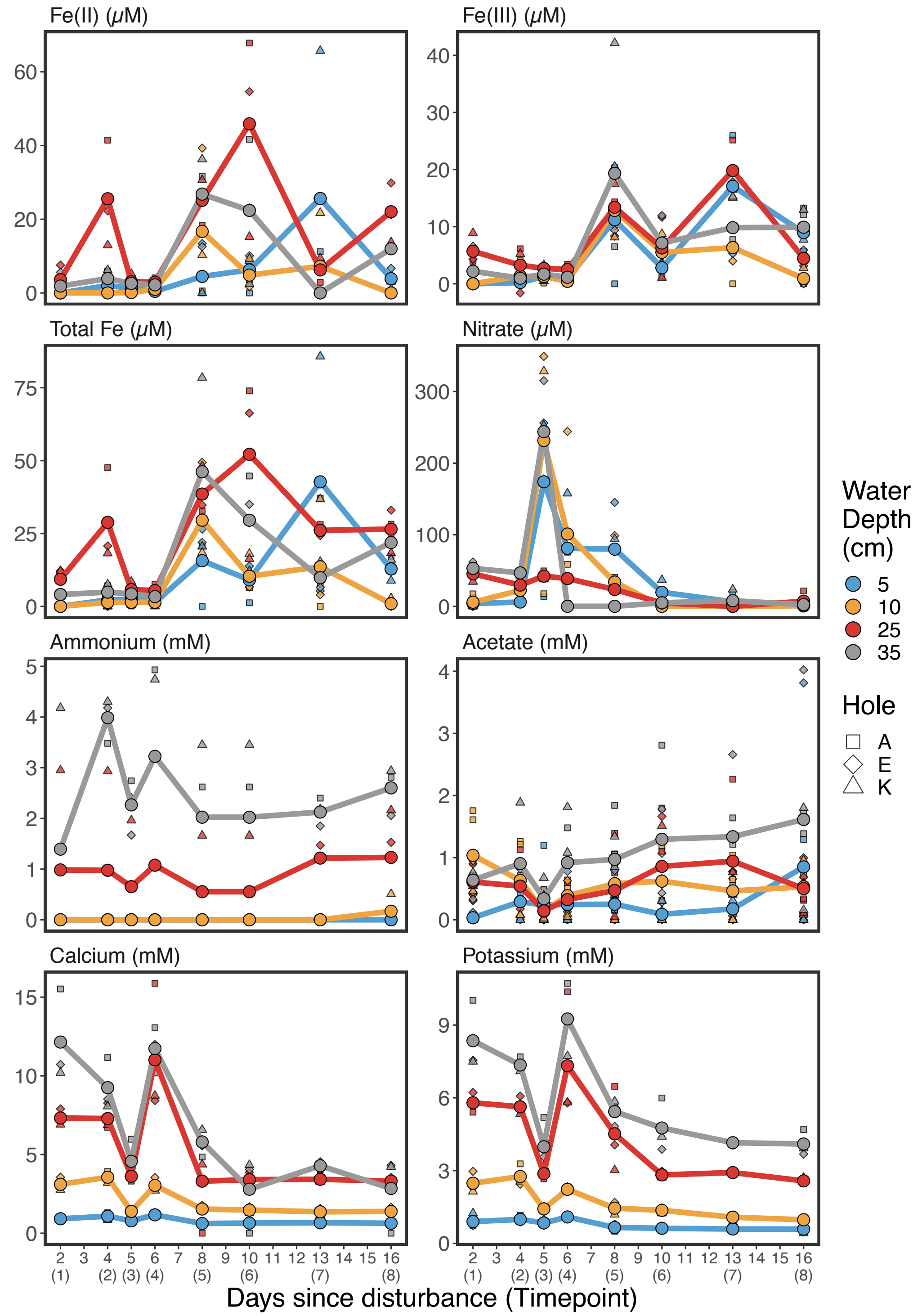

Figure S6: Physicochemistry. Iron, nitrate, ammonium, acetate, $Ca^{2+}$, and $K^{+}$ measurements. The x-axis shows days since disturbance, y-axis the respective units.

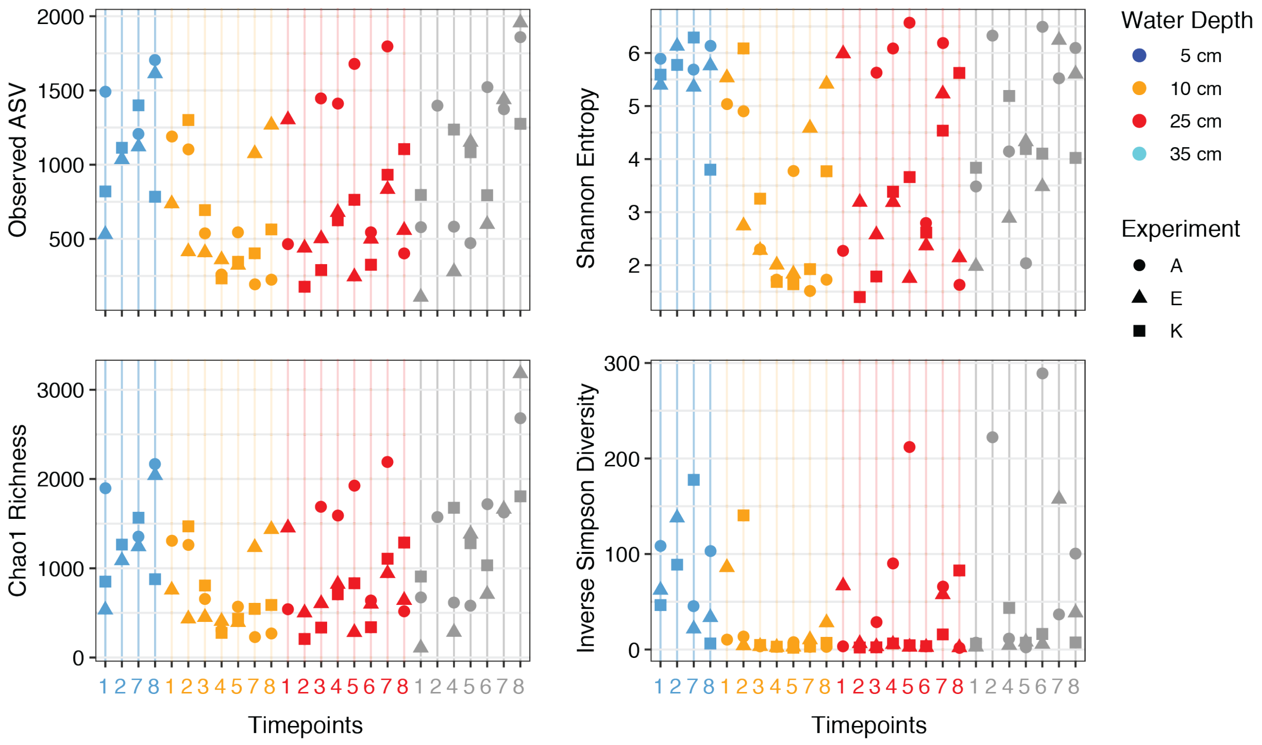

Figure S7: Individual diversity indices of all samples showing the decrease in diversity in the bloom, especially in layer 2 and 3.

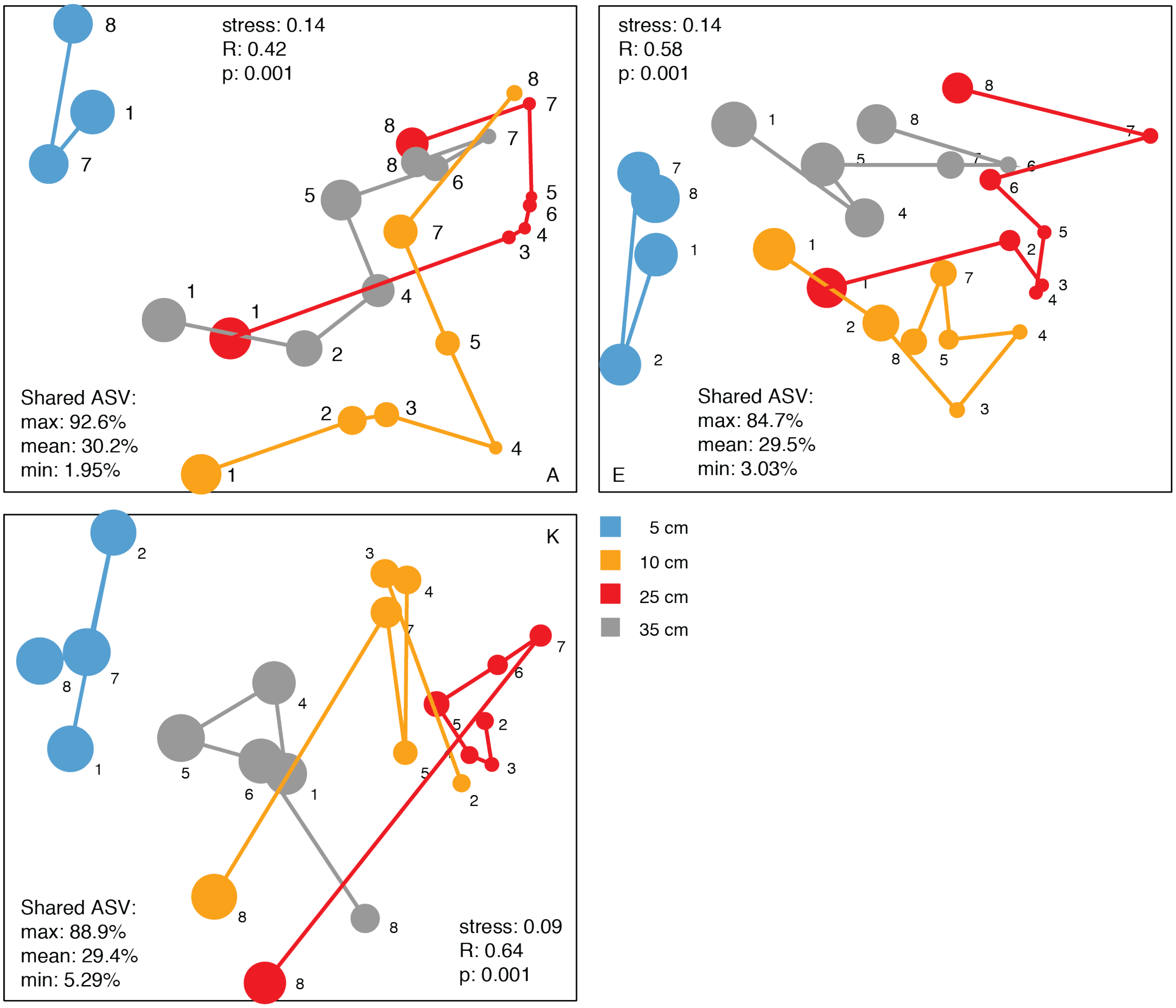

Figure S8: Trajectories of community structure in hole A, E and K. Circle size represents average Shannon Diversity across three replicate holes. Sampling time points are indicated as numbers. ASV: Amplicon Sequence Variant. Shared ASV show maximal, minimal and average percentage of shared ASV between any pair of two samples. Analysis of similarity (ANOSIM) was used to test whether the communities of each depth layer were similar. R and p values show that the groups were overlapping, but significantly different.

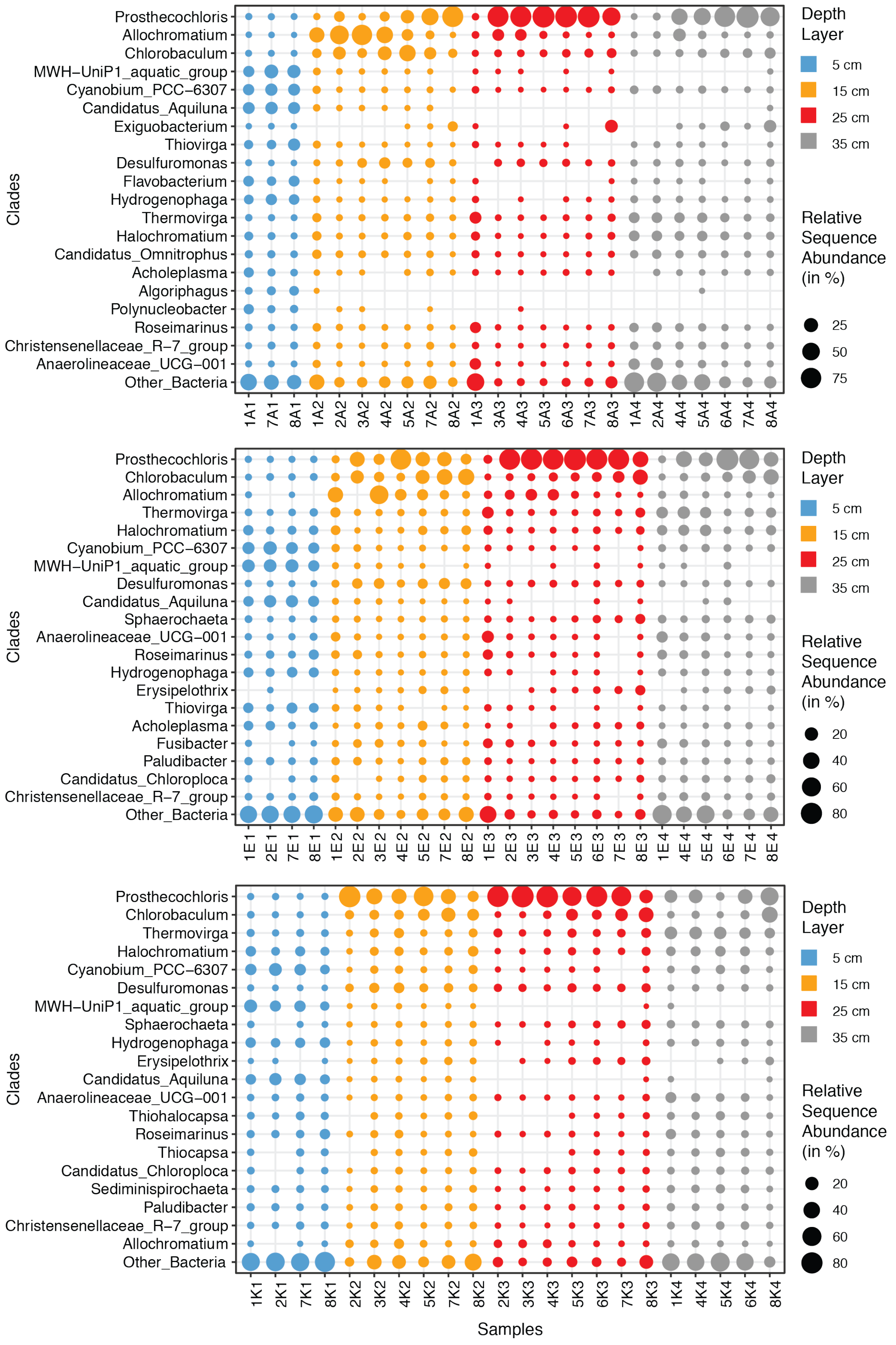

Figure S9A: Relative sequence abundance of the 20 most prominent genera in all three replicate experiments. Sample names include time point, experiment and depth, e.g. 3E2: Timepoint 3, E-hole, depth 3 (25 cm). A samples in top panel, E in middle panel, K in bottom panel).

179

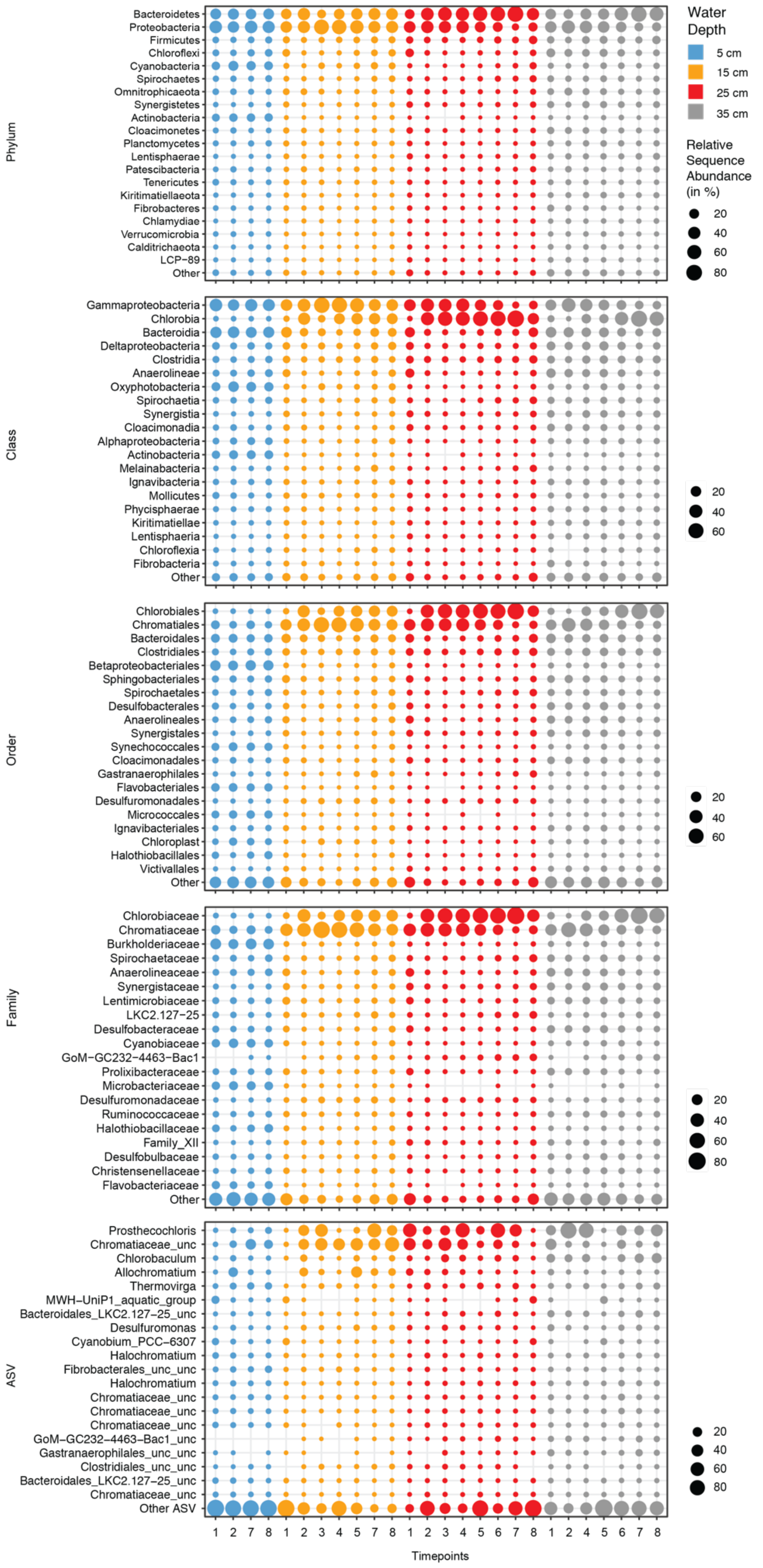

Phylum
Bacteroidetes
Proteobacteria
Firmicutes
Chloroflexi
Cyanobacteria
Spirochaetes
Omnitrophicaeota
Synergistetes
Actinobacteria
Cloacimonetes
Planctomycetes
Lentisphaerae
Patescibacteria
Tenericutes
Kiritimatiellaeota
Fibrobacteres
Chlamydiae
Verrucomicrobia
Calditrichaeota
LCP−89
Other
Class
Gammaproteobacteria
Chlorobia
Bacteroidia
Deltaproteobacteria
Clostridia
Anaerolineae
Oxyphotobacteria
Spirochaetia
Synergistia
Cloacimonadia
Alphaproteobacteria
Actinobacteria
Melainabacteria
Ignavibacteria
Mollicutes
Phycisphaerae
Kiritimatiellae
Lentisphaeria
Chloroflexia
Fibrobacteria
Other
Order
Chlorobiales
Chromatiales
Bacteroidales
Clostridiales
Betaproteobacteriales
Sphingobacteriales
Spirochaetales
Desulfobacterales
Anaerolineales
Synergistales
Synechococcales
Cloacimonadales
Gastranaerophilales
Flavobacteriales
Desulfuromonadales
Micrococcales
Ignavibacteriales
Chloroplast
Halothiobacillales
Victivallales
Other
Family
Chlorobiaceae
Chromatiaceae
Burkholderiaceae
Spirochaetaceae
Anaerolineaceae
Synergistaceae
Lentimicrobiaceae
LKC2.127−25
Desulfobacteraceae
Cyanobiaceae
GoM−GC232−4463−Bac1
Prolixibacteraceae
Microbacteriaceae
Desulfuromonadaceae
Ruminococcaceae
Halothiobacillaceae
Family_XII
Desulfobulbaceae
Christensenellaceae
Flavobacteriaceae
Other
ASV
Prosthecochloris
Chromatiaceae_unc
Chlorobaculum
Allochromatium
Thermovirga
MWH−UniP1_aquatic_group
Bacteroidales_LKC2.127−25_unc
Desulfuromonas
Cyanobium_PCC−6307
Halochromatium
Fibrobacterales_unc_unc
Halochromatium
Chromatiaceae_unc
Chromatiaceae_unc
Chromatiaceae_unc
GoM−GC232−4463−Bac1_unc
Gastranaerophilales_unc_unc
Clostridiales_unc_unc
Bacteroidales_LKC2.127−25_unc
Chromatiaceae_unc
Other ASV
1 2 7 8 1 2 3 4 5 7 8 1 2 3 4 5 6 7 8 1 2 4 5 6 7 8
Timepoints
Water Depth
5 cm
15 cm
25 cm
35 cm
Relative Sequence Abundance (in %)
20
40
60
80
20
40
60
20
40
60
20
40
60
80
20
40
60
80

Figure S9B: Relative sequence abundance of the 20 most prominent clades on phylum, class, order and family level, as well as the 20 most sequence abundant ASVs (amplicon sequence variants). The suffix _unc and _unc_unc refers to ASV of an unclassified genus or family, respectively. These ASV belong to uncultured lineages and - due to their high relative abundance - are very likely not sequence errors. Values represent average across three replicate experiments. Note: Based on the new SILVA taxonomy, the Chlorobi are now a subphylum of the Bacteroidetes.

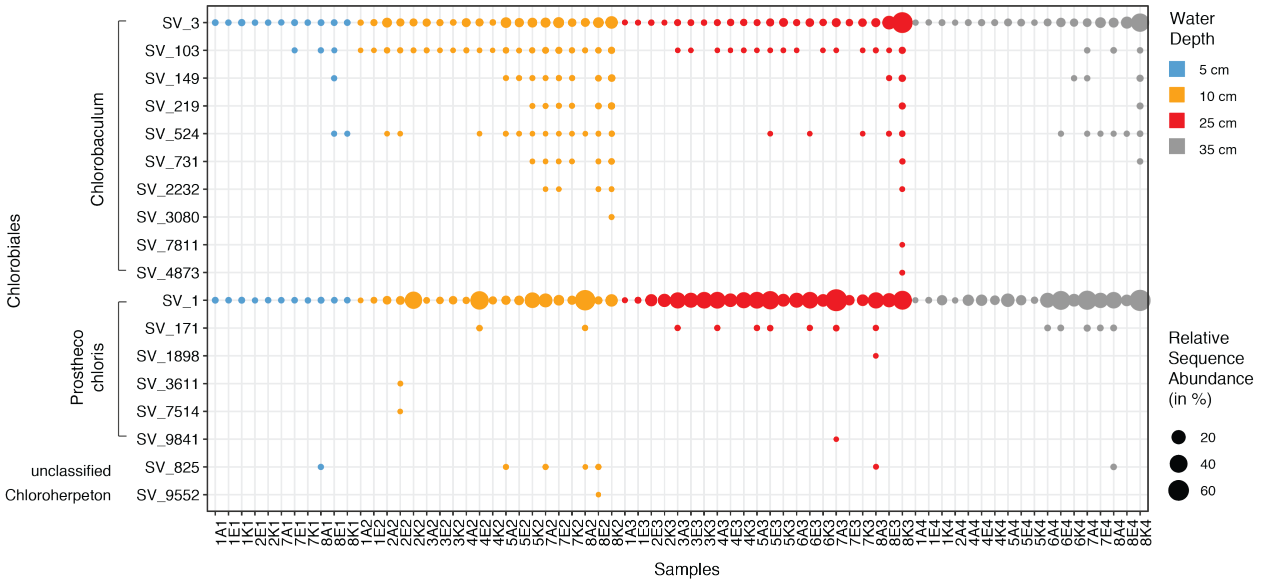

Figure S10: Relative sequence abundance of ASV within the Chlorobiales order. Sample names include time point, experiment and depth, e.g. 3E2: Timepoint 3, E-hole, depth 3 (25 cm). All samples, and replicate experiments are shown.

| | 5 cm | 10 cm | 25 cm | 35 cm |
|---|---|---|---|---|
| Prosthecochloris | 1 | 2.23 | 2.61 | 2.07 |
| Chlorobaculum | 1 | 2.37 | 2.24 | 2.45 |
| Allochromatium | 1 | 5.76 | 3.73 | 1.95 |
| Thermovirga | 1 | 1.05 | 2.05 | 1.64 |
| Halochromatium | 1 | 1.64 | 1.95 | 1.47 |
| Cyanobium_PCC-6307 | 1 | 1.03 | 0.83 | 1.04 |
| MWH-UniP1_aquatic_group | 1 | 1.15 | 0.95 | 0.99 |
| Desulfuromonas | 1 | 3.77 | 5.27 | 3.09 |
| Candidatus_Aquiluna | 1 | 0.76 | 0.56 | 0.86 |
| Hydrogenophaga | 1 | 1.49 | 1.11 | 1.06 |
| Sphaerochaeta | 1 | 1.21 | 1.57 | 1.83 |
| Thiovirga | 1 | 0.73 | 0.87 | 0.57 |
| Exiguobacterium | 1 | 3.74 | 6.61 | 1.19 |
| Anaerolineaceae_UCG-001 | 1 | 0.84 | 1.47 | 1.33 |
| Flavobacterium | 1 | 0.67 | 0.49 | 0.65 |
| Roseimarinus | 1 | 1.54 | 2.37 | 2.00 |
| Erysipelothrix | 1 | 1.25 | 1.54 | 2.32 |
| Candidatus_Chloroploca | 1 | 1.80 | 2.45 | 1.67 |
| Christensenellaceae_R-7_group | 1 | 1.46 | 2.15 | 1.37 |
| Paludibacter | 1 | 1.51 | 1.79 | 1.74 |
| Other | 1 | 1.42 | 1.59 | 1.30 |

Figure S11: Relative change of sequence abundance of ASVs between surface (V1) and deeper layers (V2-4)

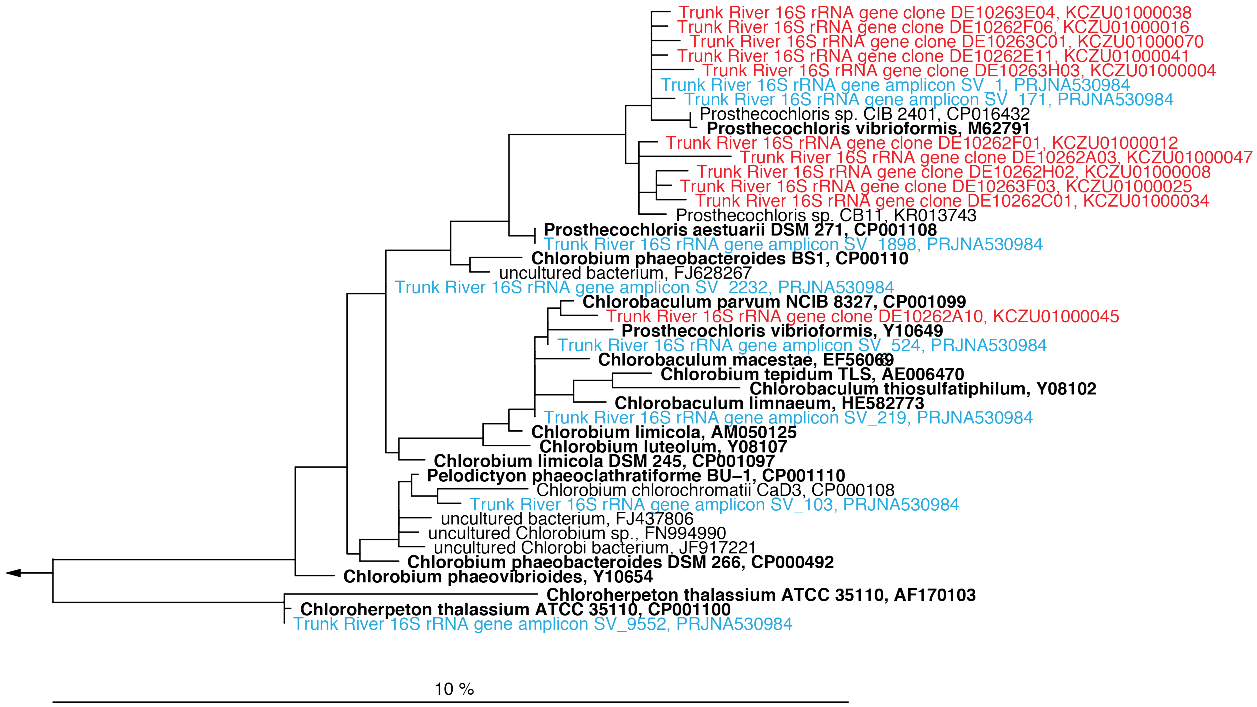

Figure S12: A phylogenetic tree depicting the affiliation of 16S rRNA amplicon sequence variants (blue), 16S rRNA gene library sequences (red) and neighboring reference sequences (cultured isolates are in bold face). The Genbank accession of provided in the node labels. Scale bar shows estimated sequence divergence. A table with nucleotide sequences of *Chlorobiales* ASVs, as well as a table of ASV frequencies across samples is publicly available under https://doi.pangaea.de/10.1594/PANGAEA.900354.

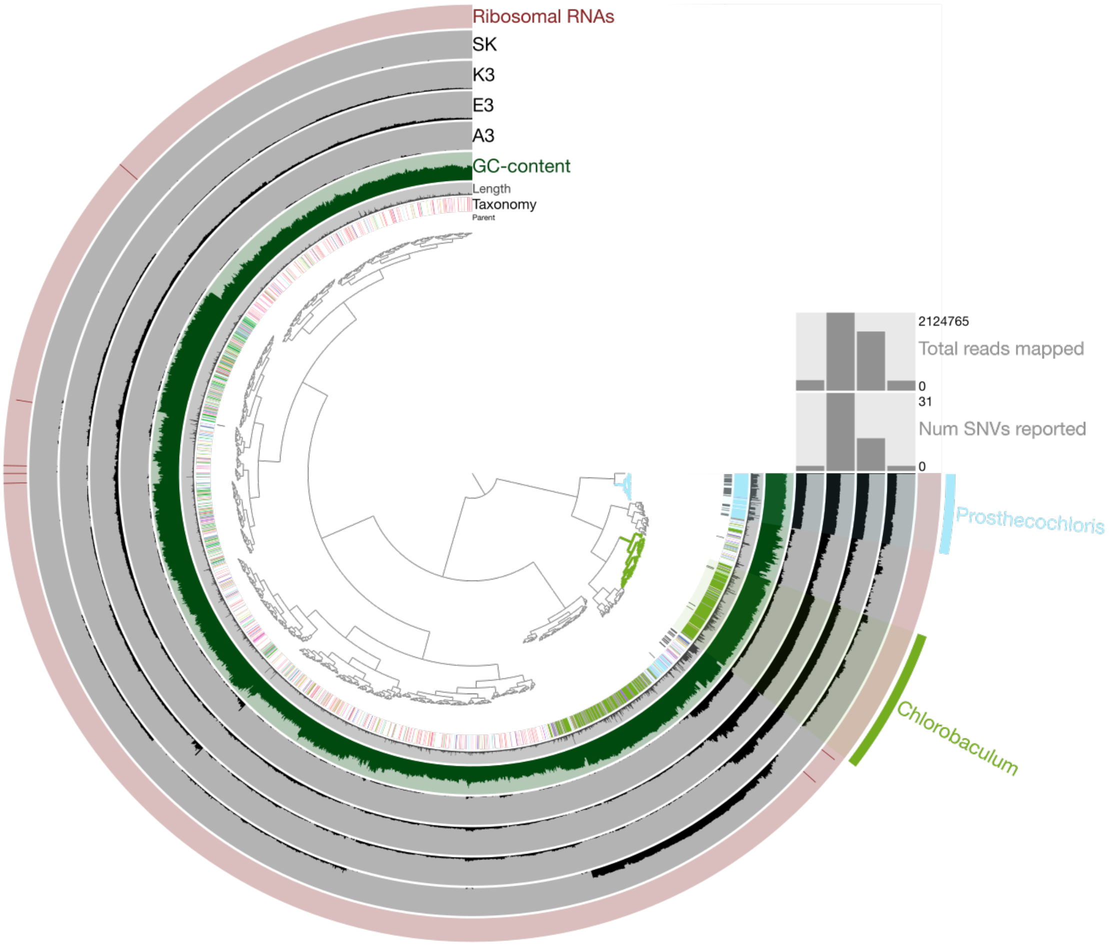

Figure S13: Circular map of metagenome-assembled genomes (MAGs) visualized using Anvi'o. The clustering dendrogram for contigs is based upon taxonomy, contig length, GC-content, and ribosomal RNA operons (from inner to outer rings) [2]. Bin 10 (Prosthecochloris sp.) and Bin 6 (Chlorobaculum sp.) metagenome-assembled genomes are highlighted.

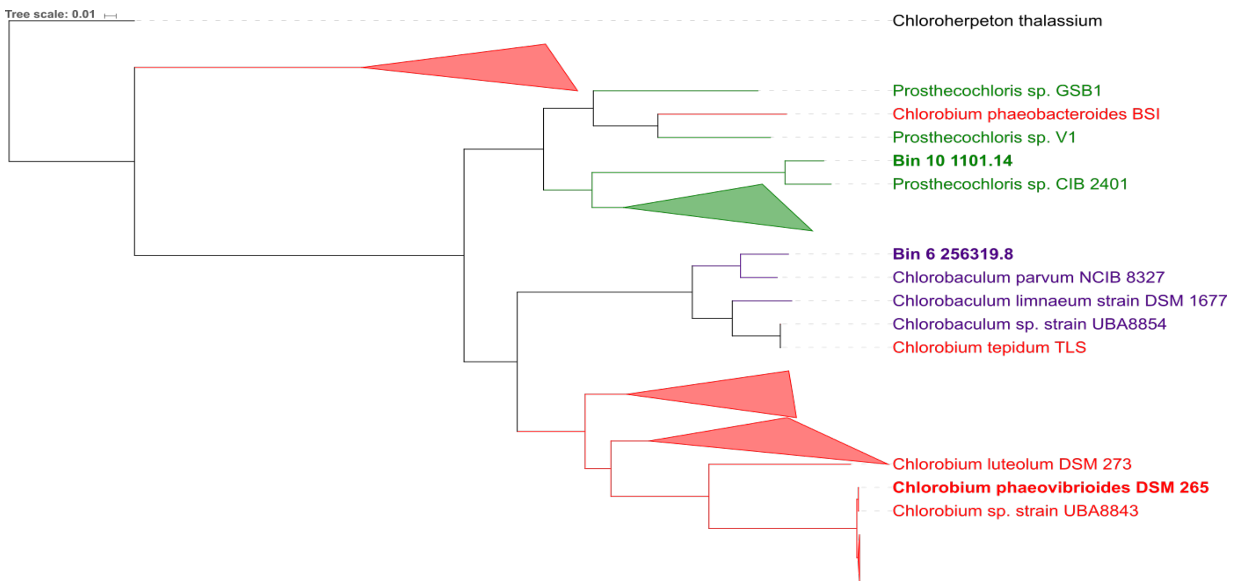

Figure S14: Hierarchical clustering analysis showing Chlorobaculum parvum NCBI 8327 as the closest genome to Bin 6 (ANI value 85.13%). Prosthecochloris sp. CIB 2401 is the closest genome to Bin 10. The scale bar represents the average number of substitutions per site.

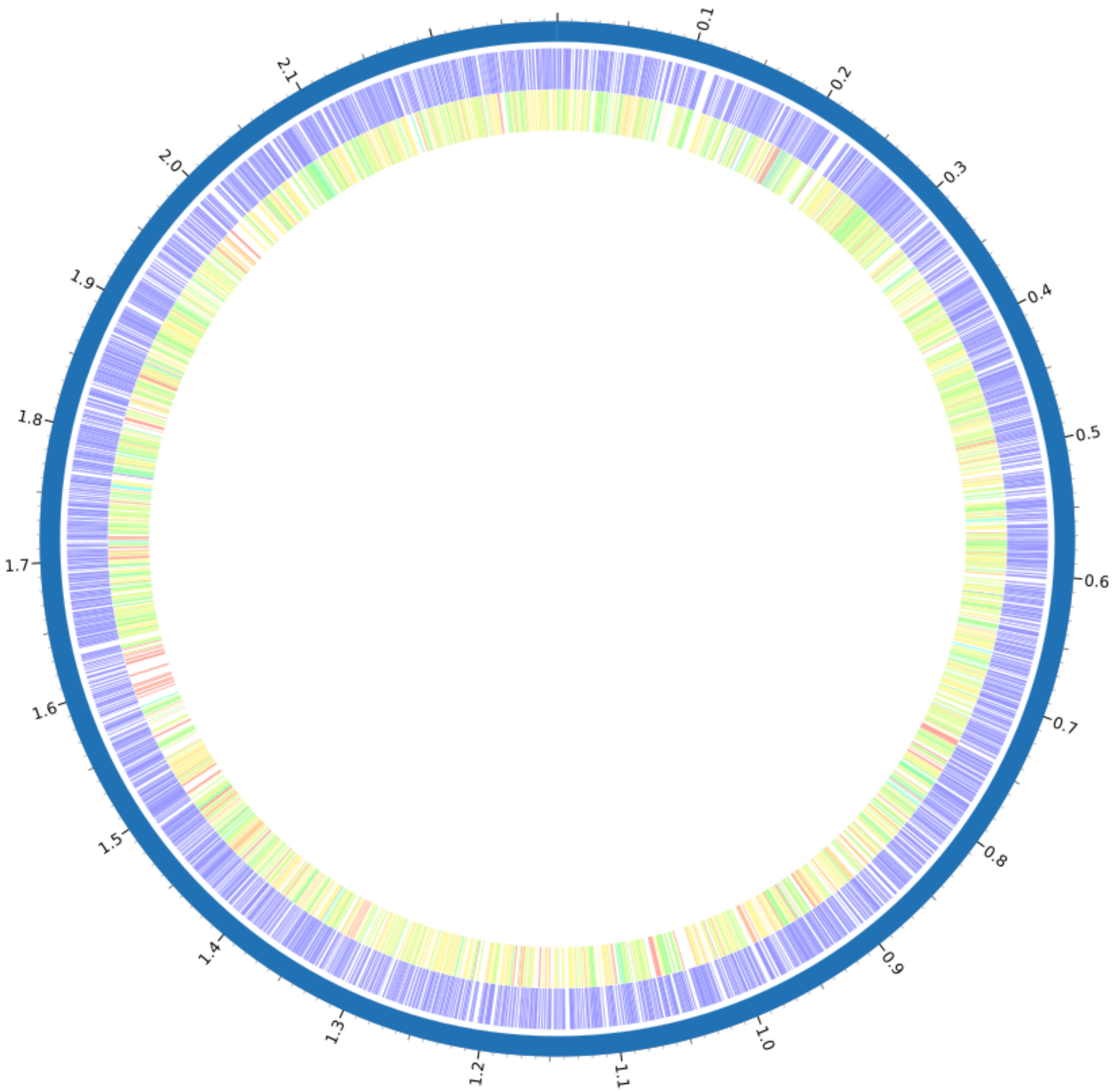

Percent protein sequence identity

| | | | | | | | | | | | | | | | |
|---|---|---|---|---|---|---|---|---|---|---|---|---|---|---|---|
| Bidirectional best hit | 100 | 99.9 | 99.8 | 99.5 | 99 | 98 | 95 | 90 | 80 | 70 | 60 | 50 | 40 | 30 | 20 | 10 |
| Unidirectional best hit | 100 | 99.9 | 99.8 | 99.5 | 99 | 98 | 95 | 90 | 80 | 70 | 60 | 50 | 40 | 30 | 20 | 10 |

Figure S15: Protein comparison of the reconstructed population genome (Bin 6) compared against the genome closest neighbor Chlorobaculum parvum NCBI 8327. Rings from outside to inside: the contig of the reference species, the reference bacterial species and the potentially novel population genome with the color scale representing the protein similarity.

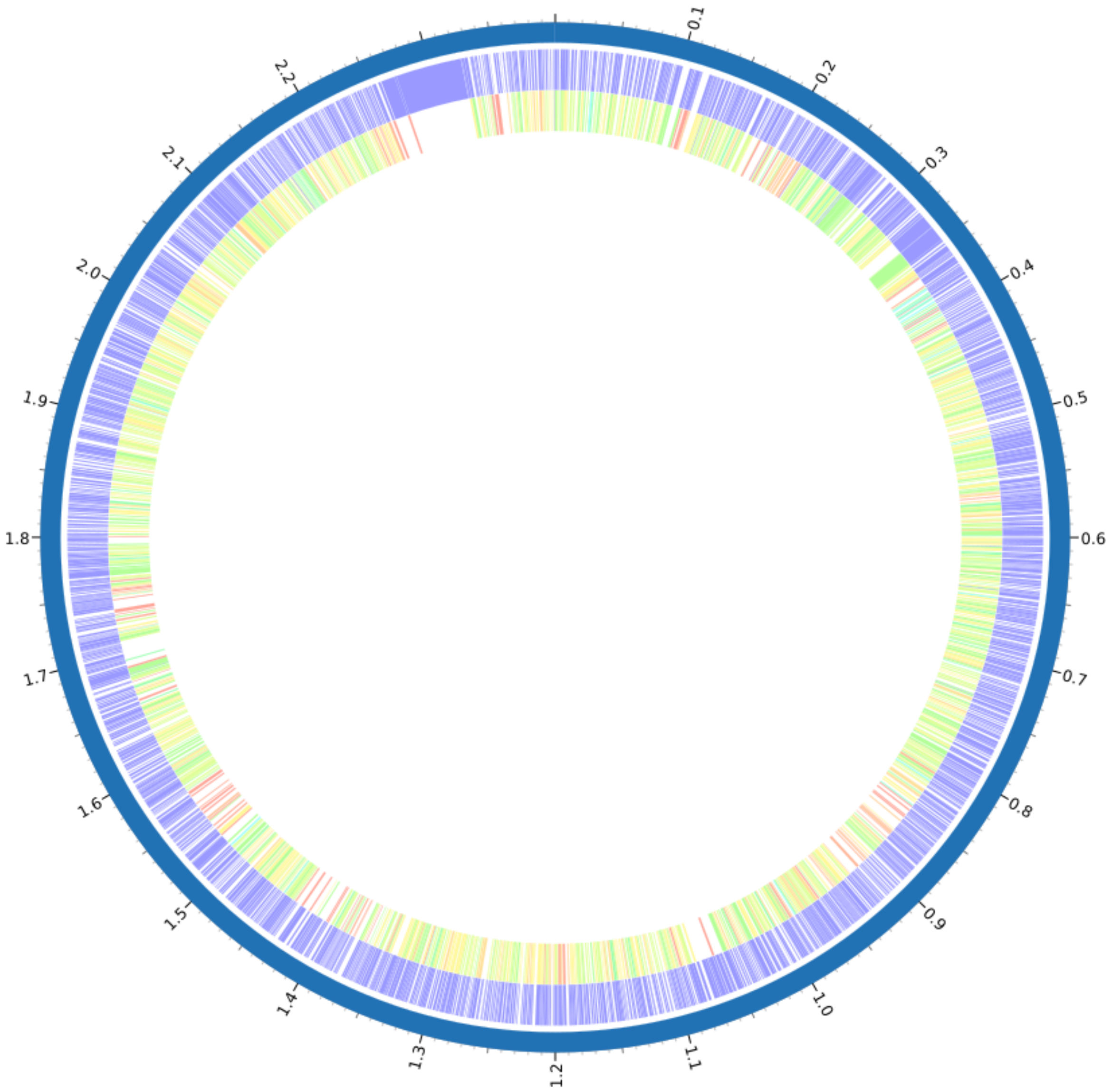

Percent protein sequence identity

| | | | | | | | | | | | | | | | | |
|---|---|---|---|---|---|---|---|---|---|---|---|---|---|---|---|---|
| Bidirectional best hit | 100 | 99.9 | 99.8 | 99.5 | 99 | 98 | 95 | 90 | 80 | 70 | 60 | 50 | 40 | 30 | 20 | 10 |
| Unidirectional best hit | 100 | 99.9 | 99.8 | 99.5 | 99 | 98 | 95 | 90 | 80 | 70 | 60 | 50 | 40 | 30 | 20 | 10 |

Figure S16: Protein comparison of the reconstructed population genome (Bin 10) compared against the genome closest neighbor Prosthecochloris sp. CIB 2401. Rings from outside to inside: the contig of the reference species, the reference bacterial species and the potentially novel population genome with the color scale representing the protein similarity.

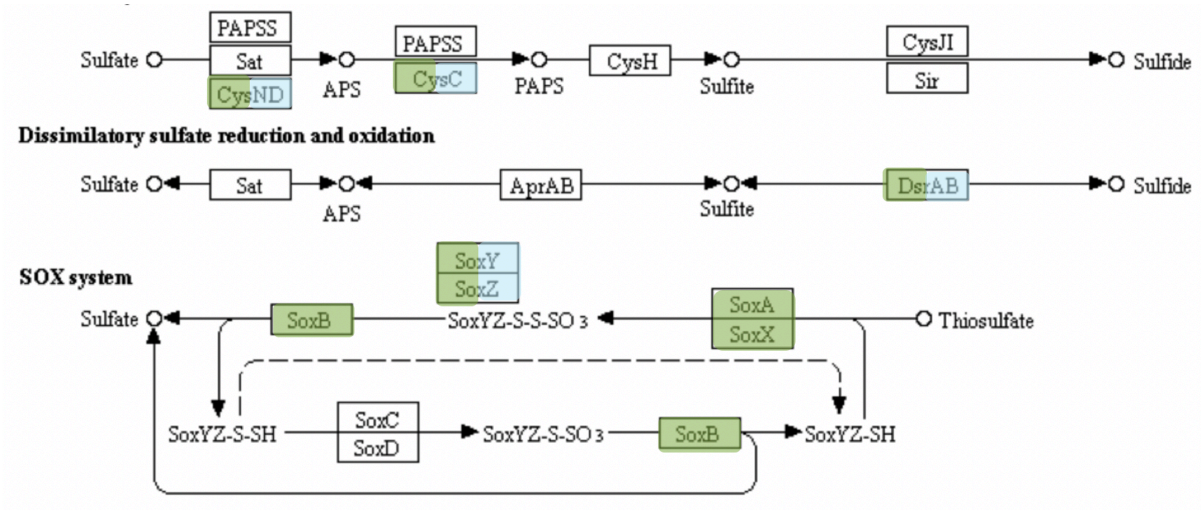

Figure 17: Genes involved in sulfur cycling

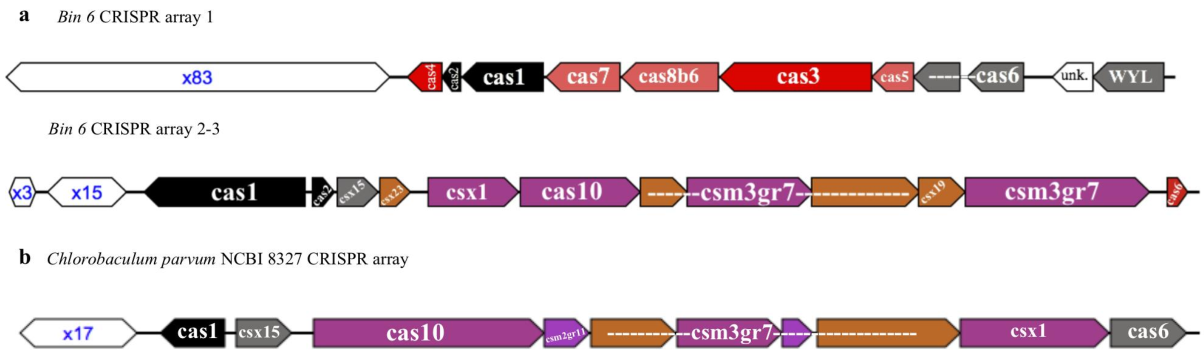

Figure S18: CRISPR arrays and cas genes predictions from (a) Bin 6 and (b) Chlorobaculum parvum NCBI 8327. CRISPR arrays are in white with the number of repeats, and cas genes are color-coded according to types/subtypes. Cas1 and cas2, which are present in most known CRISPR-Cas systems, are in black; cas3 and cas10, which are the signatures of CRISPR-Cas systems types along with Cas9 (Type II), are in red (Type I) and purple (Type III), respectively; variants of the multi-subunit complex (csm), cas10, and csx Type III genes are also in purple, and dispensable components, such as cas6, are in gray [7].

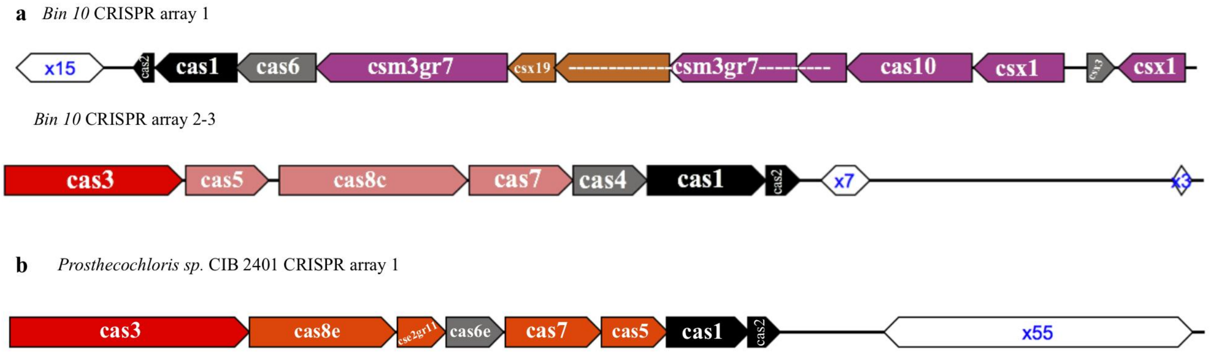

Figure S19. CRISPR arrays and cas genes predictions from (a) Bin 10 and (b) Prosthecochloris sp. CIB 2401. CRISPR arrays are in white with the number of repeats, and cas genes are color-coded according to types/subtypes. Cas1 and cas2, which are present in most known CRISPR-Cas systems, are in black; cas3 and cas10, which are the signatures of CRISPR-Cas systems types along with Cas9 (Type II), are in red (Type I) and purple (Type III), respectively; variants of the multi-subunit complex (csm), cas10, and csx Type III genes are also in purple, and dispensable components, such as cas6, are in gray [7].

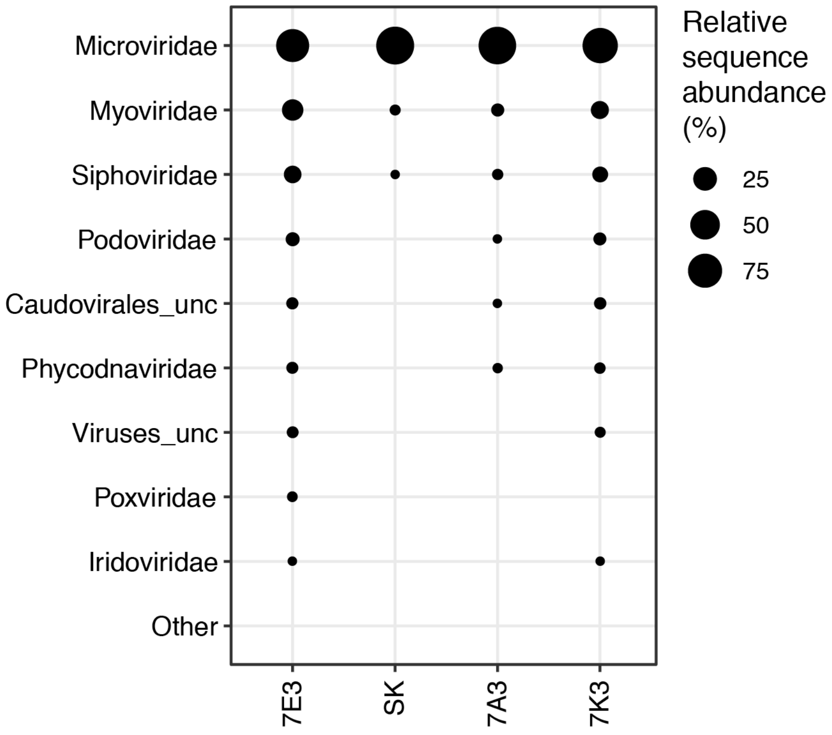

Figure S20. Relative sequence abundance of viral family-level clades based on viral sequences retrieved from metagenomes of timepoint 7 (7A3, 7E3 and 7K3) and from a phototrophic enrichment culture (SK). Unclassified family-level clades are marked by the suffix _unc.

Table S1: Overview of sequencing output and diversity indices

| | M.reads | M.sobs.r.mean | M.chao1.r.mean | M.invs.r.mean | M.SSOabs.nr | M.SSOrel.nr | M.shan.r.mean | M.SSOabs.pc | M.SSOrel.pc |
|---|---|---|---|---|---|---|---|---|---|
| 1A1 | 115879 | 1489 | 1893 | 108 | 9 | 28 | 5.9 | 0.4 | 1.3 |
| 1A2 | 13215 | 529 | 533 | 62 | 0 | 6 | 5.4 | 0 | 1.1 |
| 1A3 | 32266 | 821 | 853 | 47 | 1 | 6 | 5.6 | 0.1 | 0.7 |
| 1A4 | 30906 | 1033 | 1080 | 138 | 0 | 15 | 6.1 | 0 | 1.4 |
| 1E1 | 57122 | 1116 | 1244 | 88 | 1 | 16 | 5.8 | 0.1 | 1.2 |
| 1E2 | 63196 | 1205 | 1361 | 46 | 2 | 13 | 5.7 | 0.1 | 0.9 |
| 1E3 | 65596 | 1124 | 1247 | 22 | 2 | 12 | 5.4 | 0.2 | 0.9 |
| 1E4 | 65227 | 1410 | 1564 | 179 | 7 | 37 | 6.3 | 0.4 | 2.3 |
| 1K1 | 138540 | 1687 | 2176 | 103 | 6 | 28 | 6.1 | 0.2 | 1.2 |
| 1K4 | 125417 | 1623 | 2022 | 34 | 3 | 25 | 5.8 | 0.1 | 1.1 |
| 2A2 | 49647 | 784 | 861 | 6 | 0 | 13 | 3.8 | 0 | 1.5 |
| 2A4 | 41419 | 1180 | 1321 | 10 | 3 | 13 | 5 | 0.2 | 1 |
| 2E1 | 19251 | 738 | 754 | 86 | 0 | 5 | 5.5 | 0 | 0.7 |
| 2E2 | 53217 | 1109 | 1261 | 13 | 3 | 25 | 4.9 | 0.2 | 1.9 |
| 2E3 | 29494 | 412 | 437 | 4 | 0 | 1 | 2.7 | 0 | 0.2 |
| 2K1 | 44081 | 1301 | 1469 | 140 | 3 | 19 | 6.1 | 0.2 | 1.3 |
| 2K2 | 83011 | 538 | 652 | 3 | 0 | 4 | 2.3 | 0 | 0.6 |
| 2K3 | 46269 | 412 | 453 | 3 | 0 | 2 | 2.3 | 0 | 0.4 |
| 3A2 | 63664 | 702 | 813 | 5 | 0 | 8 | 3.3 | 0 | 1 |
| 3A3 | 57711 | 258 | 309 | 3 | 0 | 1 | 1.7 | 0 | 0.3 |
| 3E2 | 48213 | 356 | 409 | 2 | 0 | 4 | 2 | 0 | 1 |
| 3E3 | 57162 | 237 | 268 | 3 | 0 | 3 | 1.7 | 0 | 1.1 |
| 3K2 | 26817 | 542 | 570 | 8 | 0 | 3 | 3.8 | 0 | 0.5 |
| 3K3 | 63876 | 327 | 381 | 3 | 0 | 3 | 1.8 | 0 | 0.8 |
| 4A2 | 80676 | 347 | 440 | 2 | 0 | 1 | 1.6 | 0 | 0.2 |
| 4A3 | 61320 | 191 | 226 | 3 | 0 | 0 | 1.5 | 0 | 0 |
| 4A4 | 69614 | 1080 | 1224 | 10 | 1 | 10 | 4.6 | 0.1 | 0.8 |

| | | | | | | | | | |
|---|---|---|---|---|---|---|---|---|---|
| 4E2 | 145729 | 405 | 560 | 3 | 2 | 8 | 1.9 | 0.3 | 1.3 |
| 4E3 | 53079 | 224 | 262 | 3 | 0 | 1 | 1.7 | 0 | 0.4 |
| 4E4 | 68318 | 1266 | 1435 | 28 | 1 | 26 | 5.4 | 0.1 | 1.7 |
| 4K2 | 25595 | 562 | 587 | 7 | 0 | 4 | 3.8 | 0 | 0.7 |
| 4K3 | 66934 | 460 | 542 | 3 | 0 | 4 | 2.3 | 0 | 0.7 |
| 4K4 | 53867 | 1308 | 1445 | 67 | 2 | 26 | 6 | 0.1 | 1.7 |
| 5A2 | 49259 | 440 | 494 | 6 | 0 | 3 | 3.2 | 0 | 0.6 |
| 5A3 | 53923 | 178 | 214 | 2 | 0 | 0 | 1.4 | 0 | 0 |
| 5A4 | 93272 | 1443 | 1690 | 29 | 2 | 35 | 5.6 | 0.1 | 2 |
| 5E2 | 82674 | 497 | 621 | 3 | 1 | 9 | 2.6 | 0.2 | 1.4 |
| 5E3 | 70366 | 296 | 346 | 2 | 0 | 7 | 1.8 | 0 | 1.9 |
| 5E4 | 68855 | 1408 | 1590 | 92 | 5 | 15 | 6.1 | 0.3 | 0.9 |
| 5K2 | 91040 | 680 | 838 | 6 | 1 | 14 | 3.2 | 0.1 | 1.6 |
| 5K3 | 54366 | 627 | 705 | 6 | 0 | 5 | 3.4 | 0 | 0.7 |
| 5K4 | 73476 | 1679 | 1913 | 221 | 1 | 35 | 6.6 | 0 | 1.7 |
| 6A3 | 42452 | 249 | 282 | 3 | 0 | 1 | 1.8 | 0 | 0.4 |
| 6A4 | 46736 | 763 | 835 | 4 | 1 | 13 | 3.7 | 0.1 | 1.5 |
| 6E3 | 63322 | 541 | 628 | 3 | 1 | 6 | 2.8 | 0.2 | 0.9 |
| 6E4 | 71477 | 501 | 597 | 2 | 1 | 3 | 2.3 | 0.2 | 0.5 |
| 6K3 | 27235 | 325 | 340 | 4 | 0 | 2 | 2.6 | 0 | 0.6 |
| 6K4 | 123486 | 1801 | 2195 | 65 | 2 | 45 | 6.2 | 0.1 | 1.9 |
| 7A1 | 46214 | 832 | 937 | 57 | 1 | 16 | 5.2 | 0.1 | 1.7 |
| 7A2 | 93221 | 926 | 1121 | 16 | 0 | 17 | 4.5 | 0 | 1.4 |
| 7A3 | 97430 | 399 | 515 | 2 | 0 | 7 | 1.6 | 0 | 1.3 |
| 7A4 | 67784 | 556 | 643 | 2 | 0 | 6 | 2.1 | 0 | 0.9 |
| 7E1 | 78709 | 1108 | 1284 | 82 | 0 | 20 | 5.6 | 0 | 1.5 |
| 7E2 | 79334 | 576 | 670 | 7 | 0 | 11 | 3.5 | 0 | 1.5 |
| 7E3 | 11406 | 108 | 108 | 3 | 0 | 1 | 2 | 0 | 0.9 |
| 7E4 | 61491 | 794 | 899 | 6 | 1 | 17 | 3.8 | 0.1 | 1.8 |
| 7K1 | 68100 | 1399 | 1564 | 224 | 2 | 21 | 6.3 | 0.1 | 1.3 |
| 7K2 | 30559 | 578 | 607 | 11 | 1 | 8 | 4.1 | 0.2 | 1.3 |

| | | | | | | | | | |
|---|---|---|---|---|---|---|---|---|---|
| 7K3 | 23119 | 279 | 285 | 4 | 0 | 0 | 2.9 | 0 | 0 |
| 8A1 | 167167 | 1231 | 1663 | 44 | 3 | 14 | 5.2 | 0.2 | 0.7 |
| 8A2 | 89281 | 472 | 579 | 2 | 0 | 8 | 2 | 0 | 1.3 |
| 8A3 | 95105 | 1151 | 1399 | 8 | 2 | 17 | 4.3 | 0.1 | 1.1 |
| 8A4 | 85462 | 1087 | 1286 | 7 | 2 | 22 | 4.2 | 0.1 | 1.6 |
| 8E1 | 65838 | 1518 | 1709 | 294 | 3 | 23 | 6.5 | 0.2 | 1.3 |
| 8E2 | 76104 | 599 | 697 | 6 | 2 | 12 | 3.5 | 0.3 | 1.6 |
| 8E3 | 139563 | 797 | 1040 | 16 | 1 | 21 | 4.1 | 0.1 | 1.8 |
| 8E4 | 89647 | 1380 | 1627 | 37 | 1 | 27 | 5.5 | 0.1 | 1.6 |
| 8K1 | 92541 | 1439 | 1689 | 157 | 5 | 22 | 6.3 | 0.3 | 1.2 |
| 8K2 | 325158 | 1862 | 2657 | 101 | 6 | 65 | 6.1 | 0.2 | 1.9 |
| 8K3 | 573869 | 1963 | 3078 | 39 | 16 | 74 | 5.6 | 0.4 | 1.7 |
| 8K4 | 210825 | 1273 | 1784 | 8 | 3 | 39 | 4 | 0.1 | 1.8 |

Table S2: Genome statistics. Summary information for genomes binned determined by CheckM.

| Genome name | Genome size (MB) | GC (%) | Nr. of contigs | Completeness (%) | Contamination (%) | Heterogeneity (%) |
|---|---|---|---|---|---|---|
| Bin 6 | 2.46 | 56.2 | 235 | 97.84 | 0 | 0 |
| Bin 10 | 2.38 | 51.1 | 112 | 93.1 | 0 | 0 |

Table S3: Average nucleotide identity (ANI) comparisons to the closest relatives using OrthoANIu algorithm [14].

| Name | Contigs | Total length (bp) | GC content (%) | ANI (%) |
|---|---|---|---|---|
| Bin 6 | 235 | 2,460,404 | 56.35 | 85.13 |
| Chlorobaculum parvum NCIB 8327 | 1 | 2,289,249 | 55.8 | |
| Bin 10 | 112 | 2,375,588 | 51.44 | 85.82 |
| Prosthecochloris sp. CIB 2401 | 1 | 2,399,849 | 52.13 | |

Table S4: Oxidative phosphorylation and chlorophyll biosynthesis genes present (X) or absent (-) in metagenome-assembled genomes of bin 6 (*Chlorobaculum sp.*) and bin 10 (*Prosthecochloris sp.*).

| | Genes | Bin 6 | Bin 10 |
|---|---|---|---|
| Oxidative phosphorylation (terminal oxidases) | COX10 | X | - |
| | CyoE | X | - |
| | CyoD | X | - |
| | CyoC | X | - |
| | CyoB | X | - |
| | CyoA | X | - |
| | I | - | - |
| | II | - | - |
| | III | X | - |
| | CydA | X | X |

| | | | |
|---|---|---|---|
| | CydB | X | X |
| (Bacterio)chlorophyll biosynthesis | BchK | X | X |
| | bciD | - | - |
| | bchU | X | X |
| | bchE | X | X |
| | bchM | X | X |

Table S5: CRISPR-Cas system information for each metagenome-assembled genome

| Name | CRISPR array | contig | location (bp) | length | # spacers | DR length |
|---|---|---|---|---|---|---|
| Bin 6 | CRISPR1 | 26 | 20933-26430 | 42829 | 82 | 30 |
| Bin 6 | CRISPR2 | 145 | 613-1685 | 16001 | 14 | 35 |
| Bin 6 | CRISPR3 | 145 | 198-278 | 16001 | 2 | 35 |
| Bin 10 | CRISPR1 | 20 | 3561-4695 | 55534 | 14 | 37 |
| Bin 10 | CRISPR2 | 316 | 7079-7510 | 10315 | 6 | 32 |
| Bin 10 | CRISPR3 | 316 | 10128-10293 | 10315 | 2 | 33 |
